# Loss of Brg1 promotes seizure development via GABAergic system disruption

**DOI:** 10.64898/2026.02.25.706532

**Authors:** Roberto Pagano, Karim Abu Nahia, Amber Declève, Dorota Stadnik, Justyna Zmorzyńska, Remigiusz A. Serwa, Daniëlle Copmans, Jacek Jaworski

## Abstract

The Brahma-related gene 1 (*BRG1*) encodes the catalytic subunit of the SWItch/Sucrose Non-Fermentable (SWI/SNF) chromatin remodeling complex and plays an important role in brain development. Variants in SWI/SNF components are often found in patients with epilepsy, but it is still unclear how loss of BRG1 function contributes to seizure development. In this study, we analyzed the role of Brg1 in seizure susceptibility using zebrafish models with both pharmacological inhibition or genetic reduction of Brg1. Reduced Brg1 function caused seizure-like behavior and increased neuronal activity in larvae, while basic locomotor activity was preserved. Further analyses showed reduced expression of several GABAergic system markers. In contrast, glutamatergic markers did not show major changes. These results point to a selective impairment of inhibitory signaling. When GABA levels were increased pharmacologically, seizure-like behavior was reduced. This suggests that loss of inhibitory transmission plays an important role in the observed hyperexcitability. Unbiased omics analyses also identified changes in proteins associated with vitamin B6 binding. Treatment with active vitamin B6 reduced seizure-like behavior in larvae with reduced Brg1 function. Taken together, these results indicate that Brg1 is required for proper inhibitory neurotransmission and that partial loss of Brg1 function increases seizure susceptibility. These findings may help to better understand why mutations in chromatin remodeling genes are often associated with epilepsy and could support future studies on targeted modulation of inhibitory signaling in these conditions.

**Significance Statement:** Chromatin remodeling genes are often mutated in patients with epilepsy, but it is still unclear how these mutations lead to seizures. In this study, we show that reduced function of the chromatin remodeler Brg1 affects inhibitory neurotransmission by impairing the GABAergic system. This leads to increased neuronal activity and seizure-like behavior. Our results identify the chromatin remodeler Brg1 as an important regulator of inhibitory neurotransmission and seizure susceptibility, which may be important for understanding epilepsy associated with neurodevelopmental disorders.

## Introduction

Epilepsy is a neurological disease characterized by recurrent episodes of abnormal synchronous brain activity (seizures), which can affect consciousness or induce involuntary movements (1–3). It is a complex disease with multiple risk factors, such as genetic, structural, infectious, metabolic, immune, and neurodegenerative (2, 4). Recent research has increasingly focused on the role of epigenetic mechanisms in epileptogenesis, highlighting how chromatin remodeling affects gene expression patterns that may contribute to seizure susceptibility and epilepsy development (5, 6).

Among the key regulators of chromatin, the SWItch/Sucrose Non-Fermentable (SWI/SNF) complex has emerged as a crucial player in neurological development and disease (7, 8). It contains a catalytic core subunit, either Brahma-related gene 1 (BRG1, also known as SMARCA4) or Brahma (BRM), plus 10 to 15 Brg1/BRM-associated factors (BAFs) whose composition varies by cell type (9–12). Using ATP, this complex remodels nucleosomes to regulate transcriptional access to specific genomic regions (7, 8, 11, 13–17).

Mutations in SWI/SNF components, including *BRG1*, are linked to neurodevelopmental disorders such as Coffin–Siris syndrome, Nicolaides–Baraitser syndrome, autism spectrum disorders (ASD), schizophrenia, and epilepsy syndromes (7, 13, 18, 19). Notably, up to 26% of Coffin–Siris patients experience epileptic episodes (20–24), pointing to a connection between SWI/SNF dysfunction and epilepsy, although the underlying mechanisms remain unclear.

BRG1 and BRM ATPases share sequence similarity, but they serve distinct functions. Brg1 is essential for early development, as its absence results in embryonic lethality before implantation (25–27). Deleting *Brg1* in early postnatal hippocampal neurons reduces dendritic spine density and maturity, disrupts synaptic function, and leads to neurological abnormalities in developing mice (28). Also, Brg1 can recruit either transcriptional activators (e.g., Calcium Responsive Transactivator [CREST]–CREB Binding Protein [CBP]) or repressors (e.g., Mammalian Sin3 homolog A [mSin3A]/Histone deacetylase 1 [HDAC1]/Retinoblastoma protein [Rb]) to the promoters and enhancers of activity-related genes such as *c-Fos* and *Grin2b* (11, 14, 29). These mechanisms help explain why Brg1 mutations are associated with intellectual disability and ASD (28, 30–32); however, if and how loss of Brg1 function leads to epilepsy, a common comorbid phenotype remains to be demonstrated. In particular, it remains unclear whether the epileptic phenotype associated with Brg1 mutations reflects widespread changes in neural circuit properties due to developmental morphological changes or more selective impairments of defined types of synaptic transmission.

In this study, we aimed to investigate the role of the BAF complex, with a particular emphasis on Brg1, in seizure susceptibility using a zebrafish model. Therefore, by combining pharmacological inhibition of the BAF complex, *in vivo* CRISPR/Cas9-mediated Brg1 editing, RNA sequencing, mass spectrometry, calcium imaging, and behavioral analyses, we examined the effects of Brg1 downregulation on seizure-related phenotypes, neuronal activity, and excitatory–inhibitory transmission imbalance.

## Results

### BAF complex inhibition induces seizure-like behavior in zebrafish larvae

Mutations in the SWI/SNF chromatin remodeling complex have been linked to multiple neurodevelopmental disorders, including seizure syndromes (7, 13, 18, 19). Here, to model functional consequences of loss of BAF complex activity, we examined whether pharmacological inhibition of the BAF complex induces seizure-like activity in zebrafish. The chosen Brg1/BRM inhibitor (BAF inhibitor) was extensively utilized across various studies to elucidate the roles of chromatin remodeling in diverse biological processes, including inflammation, cancer, gene regulation, and immune responses (33–35).

Before conducting behavioral assays, the efficacy of the BAF inhibitor was validated. Embryos at 5 hours post-fertilization (hpf) were exposed overnight to concentrations ranging from 1 nM to 100 µM, and were compared with DMSO-treated larvae used as control. 10 µM was the highest concentration preserving normal morphology (**SI Appendix Fig. S1A**). In 96 hpf larvae, overnight exposure to 10 µM inhibitor reduced expression of Brg1 targets *klf2a* and *nos1* (36), confirming effective BAF inhibition (**SI Appendix Fig. S1B**).

To assess seizure development, two behavioral proxies were measured: spontaneous tail flick and body coiling (STFBC), and locomotor activity (37). Embryos were treated with the inhibitor or DMSO from 5 to 24 hpf and analyzed for STFBC (**Fig. 1A i, ii**), a behavior associated with seizure activity (37, 38). BAF inhibition (**Fig. 1A iii**) significantly increased the proportion of double-coil events that depend on glutamatergic modulation (**Fig. 1A iii**), suggesting changes in excitatory transmission in 24 hpf zebrafish (38).

**Figure 1.**
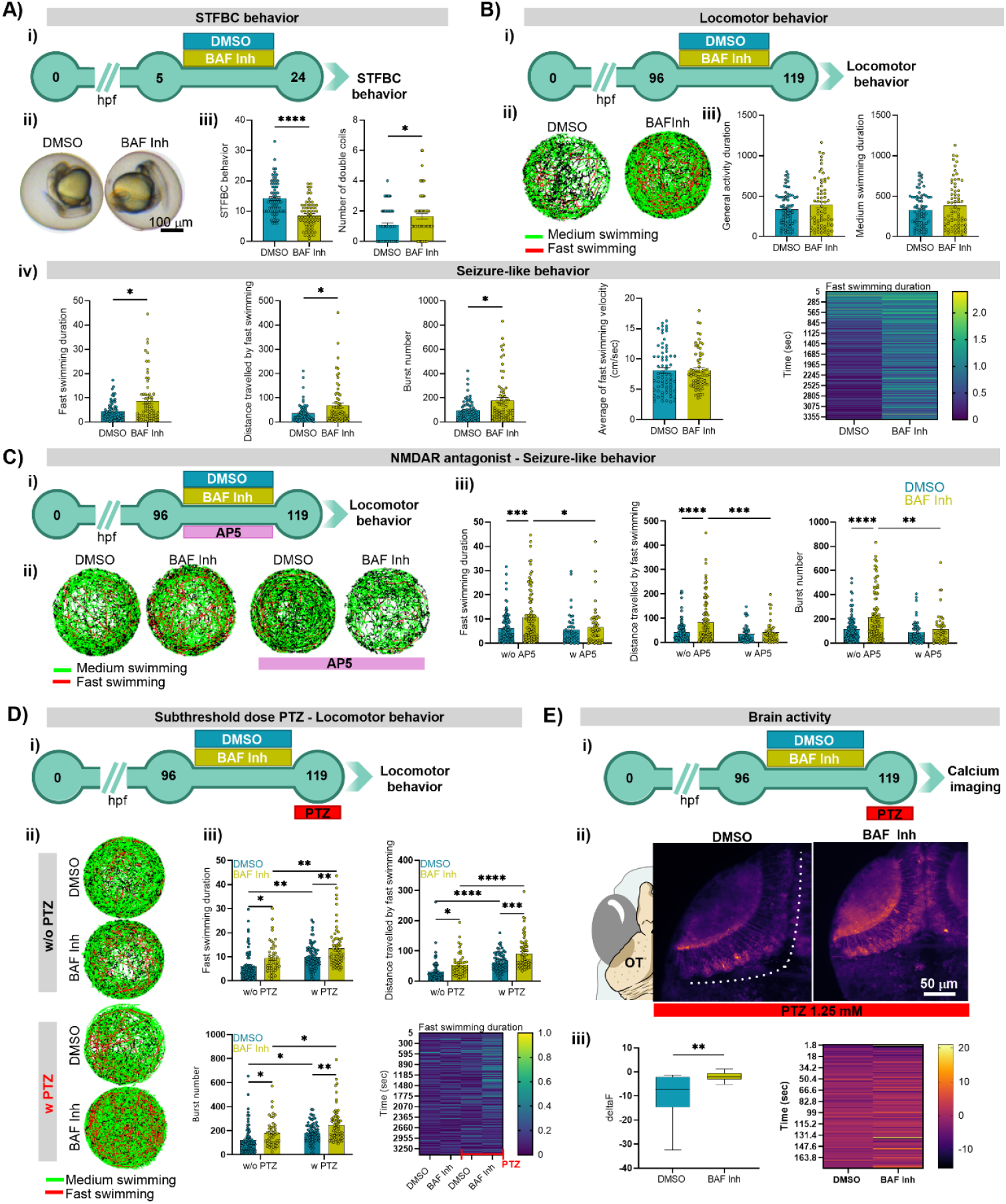
BAF complex inhibition induces seizure-like behavior in zebrafish larvae. **(A) i)** Experimental timeline. Zebrafish embryos were treated at 5 hours post-fertilization (hpf) with either DMSO or a BAF complex inhibitor (BAF Inh, 10 µM) overnight. The following day, spontaneous tail flick and coil (STFBC) behavior was assessed. **ii)** Representative images of zebrafish larvae at 24 hpf during the recording of spontaneous tail flick and body coil (STFBC) behavior following DMSO or BAF Inh. **iii)** Number of total coils (Mann-Whitney test, U = 1029, *p* < 0.0001) and double coils (Mann-Whitney test, U = 1838, *p* = 0.017) in BAF Inh-treated larvae compared to DMSO controls (n_DMSO_ = 72, n_BAFInh_ = 66). Data are presented as mean ± standard error of the mean (SEM). **(B) i)** Experimental timeline. Zebrafish larvae were treated at 96 hpf with either DMSO or a BAF Inh (10 µM) overnight. Locomotor behavior was then assessed. **ii)** Representative locomotor tracks of larvae treated with DMSO or BAF Inh. Medium swimming track (0.5-2 cm/sec) is shown in green, while fast swimming track (>2 cm/sec) is shown in red. **iii)** General locomotor activity: Duration of general activity (Mann-Whitney test, U = 2279, *p* = 0.5699) and duration of medium swimming (Mann-Whitney test, U = 2287, *p* = 0.5944) after BAF inhibitor. Data are presented as mean ± SEM. **iv)** Seizure-like behavior parameters: Fast swimming duration (Mann-Whitney test, U = 1832, *p* = 0.0139), distance traveled with fast swimming (Mann-Whitney test, U = 1865, *p* = 0.0204), number of burst episodes (Mann-Whitney test, U = 1902, *p* = 0.0307), average velocity during fast swimming (Mann-Whitney test, U = 2293, *p* = 0.6105), and a heatmap illustrating fast swimming duration over time (n_DMSO_ = 71, n_BAFInh_ = 68). Data are presented as mean ± SEM. **(C) i)** Experimental timeline. Zebrafish larvae were treated at 96 hpf with either DMSO or a BAF Inh overnight with D-2-amino-5-phosphonovalerate (AP5, 100 µM). **ii)** Representative locomotor tracks of larvae treated with DMSO or BAF Inh. Medium swimming track (0.5-2 cm/sec) is shown in green, while fast swimming track (>2 cm/sec) is shown in red. **iii)** Seizure-like behavior parameters: Fast swimming duration (two-way ANOVA: AP5 effect, F (1, 294) = 5.023, *p* < 0.00258; treatment effect, F (1, 294) = 8.387, *p* = 0.0041; Tukey post hoc test), distance traveled with fast swimming (two-way ANOVA: AP5 effect, F (1, 294) = 10.92, *p* = 0.0011; treatment effect, F (1, 294) = 10.46, *p* = 0.0014; AP5 vs. treatment interaction, F (1, 294) = 5.031, *p* = 0.0256; Tukey post hoc test), number of burst episodes (two-way ANOVA: AP5 effect, F (1, 294) = 11.24, *p* = 0.0009; treatment effect, F (1, 294) = 10.71, *p* = 0.0012; Tukey post hoc test), (n_DMSO_ = 104, n_BAFInh_ = 99, n_DMSO_AP5_ = 48, n_BAFInh_AP5_ = 47). Data are presented as mean ± SEM. **(D) i)** Experimental timeline. Zebrafish larvae were treated at 96 hpf with either DMSO or BAF Inh overnight. Locomotor behavior was assessed after a 3-minute incubation with a subthreshold dose of pentylenetetrazole (PTZ, 1.25 mM). **ii)** Representative locomotor tracks of larvae treated with DMSO or BAF Inh, with and without PTZ. **iii)** Quantification of seizure-like behavior parameters: Fast swimming duration (two-way ANOVA: PTZ effect, F(1, 259) = 22.34, *p* < 0.0001; treatment effect, F(1, 63) = 22.85, *p* < 0.0001; Tukey post hoc test), distance traveled during fast swimming (two-way ANOVA: PTZ effect, F(1, 259) = 45.24, *p* < 0.0001; treatment effect, F(1, 259) = 26.09, *p* < 0.0001; Tukey post hoc test), number of burst episodes (two-way ANOVA: PTZ effect, F(1, 259) = 19.10, *p* < 0.0001; treatment effect, F(1, 259) = 17.63, *p* < 0.0001; Tukey post hoc test) and heatmap showing the duration of fast swimming over time (n_DMSO_ = 75, n_DMSO_PTZ_ = 72, n_BAFInh_ = 47, n_BAFInh_PTZ_ = 69). Data are presented as mean ± SEM. **(E) i)** Experimental timeline. Zebrafish larvae were treated at 96 hpf with either DMSO or BAF Inh overnight. Calcium imaging was performed following a 3-minute incubation with a subthreshold dose of PTZ (1.25 mM). **ii)** Representative Z-projection (maximum intensity) images showing neuronal activity in the optic tectum (OT) of larvae treated with DMSO or BAF Inh. **iii)** Quantification of neuronal activity (Mann-Whitney test, U = 63, *p* = 0.0025) and heatmap showing temporal dynamics of neuronal activity (n_DMSO_PTZ_ = 14, n_BAFInh_PTZ_ = 22). Data are presented as Tukey box-and-whisker plots. \**p* < 0.05, \*\**p* < 0.01, \*\*\**p* < 0.001, \*\*\*\**p* < 0.0001.

To evaluate seizure-like behavior at later stages, larvae were treated from 96 to 119 hpf and monitored for one hour in the Zebrabox tracking system (**Fig. 1B i**). Swim trajectories were classified as “medium” (normal locomotion) or “fast” (seizure-like) swims (**Fig. 1B ii**) (37). Total activity and medium swim duration were unchanged, indicating intact baseline motor function (**Fig. 1B iii**). In contrast, BAF inhibition markedly increased time and distance spent in fast swimming, as well as burst frequency, without altering burst peak velocity (**Fig. 1B iv**). These results show that BAF inhibition enhances seizure-like locomotor behavior.

Given that seizures often result from an impaired excitatory/inhibitory transmission balance (39– 41) we tested whether blocking glutamatergic transmission could rescue this phenotype. Co-treatment with the N-methyl-D-aspartate receptor (NMDAR) antagonist D-2-amino-5-phosphonovalerate (AP5) abolished BAF inhibitor-induced increases in fast swim duration, distance, and burst number (**Fig. 1C i–iii**), indicating that excessive excitatory signaling mediates the seizure-like activity caused by BAF inhibition.

Next, we assessed whether prolonged BAF complex inhibition would worsen the seizures by treating larvae with the BAF inhibitor from 72 to 119 hpf (**SI Appendix Fig. S2A i**). However, extended exposure reduced overall locomotor activity and seizure-like behavior (**SI Appendix Fig. S2A ii–iv**). Most of the BAF inhibitor-treated larvae developed an opaque brain, a feature linked to neuronal death (42) (**SI Appendix Fig. S2A v**). To confirm this, we used acridine orange staining to assess cell death in live larvae treated with the inhibitor (**SI Appendix Fig. S2B i**). While no cell death was detected at 24 hours between groups, robust apoptosis emerged by 48 hours, affecting the pallium, cerebellum, and most prominently the optic tectum (OT) (**SI Appendix Fig. S2B ii–v**). Also, pro-inflammatory cytokines were upregulated after 24 hours of treatment, and inflammatory genes after 48 hours (**SI Appendix Fig. S2C i, ii**), consistent with inflammation observed in human temporal lobe epilepsy (hTLE) (42, 43). Moreover, microglial (*mpeg1*) and astroglial (*gfap*) activation increased after 24 hours of treatment (**SI Appendix Fig. S2D ii, iii**), suggesting a glial response during the initial stages of inflammation.

Overall, BAF inhibition increased double-coil events and enhanced seizure-like behavior, while also inducing pro-inflammatory responses in the brain and activating microglia and astroglia. Prolonged inhibition reduced overall locomotor activity, and was associated with brain inflammation and cell death, features often occurring with enhanced neuronal activity leading to excitotoxicity (44–46).

### BAF inhibition increases seizure-like behavior and neuronal activity following subthreshold PTZ exposure

To further confirm whether inhibition of the BAF complex induces seizure development, the GABA receptor antagonist pentylenetetrazole (PTZ), which is known to evoke clonic-like seizures in fish (47), was used. A subthreshold dose of PTZ was applied three minutes before the behavioral assay to disinhibit the system without inducing seizures on its own, allowing us to assess whether it modulated the seizure-like behavior induced by BAF inhibition (**Fig. 1D i**). BAF inhibitor treatment significantly increased the duration and distance of fast swimming, as well as the average burst number and fast swimming velocity, in both PTZ and non-PTZ conditions compared to the respective DMSO controls (**Fig. 1D ii, iii**). Notably, BAF inhibitor treated larvae exposed to PTZ exhibited the most pronounced hyperlocomotor response, indicating heightened sensitivity to subthreshold seizure-inducing stimuli (**Fig. 1D ii, iii**).

We next investigated whether this increased seizure-like behavior was accompanied by elevated brain activity. Using *Tg(HuC:GCaMP5G)* zebrafish larvae, time-lapse calcium imaging of neuronal activity was performed following three minutes of subthreshold PTZ exposure (**Fig. 1E i**). The analysis focused on the OT, as the largest effect was observed in this structure in the experiments described in the previous section (**SI Appendix Fig. S2B ii–v**). Fluorescence changes over time in the OT were used to quantify neural activity in the BAF inhibitor compared to DMSO (**Fig. 1E ii**). Analysis revealed a significant increase in calcium activity in the BAF inhibitor group compared to DMSO (**Fig. 1E iii**).

To further support our conclusion that BAF complex inhibition alters neural excitability, non-invasive local field potentials (LFPs) were recorded from the OT of larvae treated with DMSO or BAF inhibitor, followed by subthreshold PTZ exposure (**SI Appendix Fig. S3A i**). Power spectral density (PSD) analysis of the 20–140 Hz frequency range showed no significant difference in overall power between groups (**SI Appendix Fig. S3A ii, iii**). However, frequency-resolved analysis across this range revealed a broad non-significant increase in normalized PSD values in the BAF inhibitor group, starting from 40 Hz (**SI Appendix Fig. S3A iv**), suggesting enhanced PSD, including high-frequency oscillatory activity (80–140 Hz). Further examination of specific frequency bands showed no significant differences in PSD in the 30–80 Hz band (gamma waves), whereas an increasing tendency was observed in the 80–140 Hz range (high-frequency oscillations) in the BAF inhibitor group (**SI Appendix Fig. S3A v**).

These findings demonstrate that BAF complex inhibition lowers the threshold for seizure induction and promotes the hyperexcitability of the OT.

### BAF complex inhibition modulates visual epilepsy- and GABAergic-related gene expression

To characterize the transcriptional impact of the BAF complex in the brain and identify molecular pathways linking its dysfunction to seizure development, we performed RNA sequencing (RNAseq) on heads dissected after 24 hours BAF inhibition or DMSO treatment (**Fig. 2A; SI Appendix Fig. S4**). Comparison identified 1754 differentially expressed genes (DEGs) in BAF inhibitor treated larvae, of which 816 were upregulated and 938 downregulated (**Fig. 2A**).

**Figure 2.**
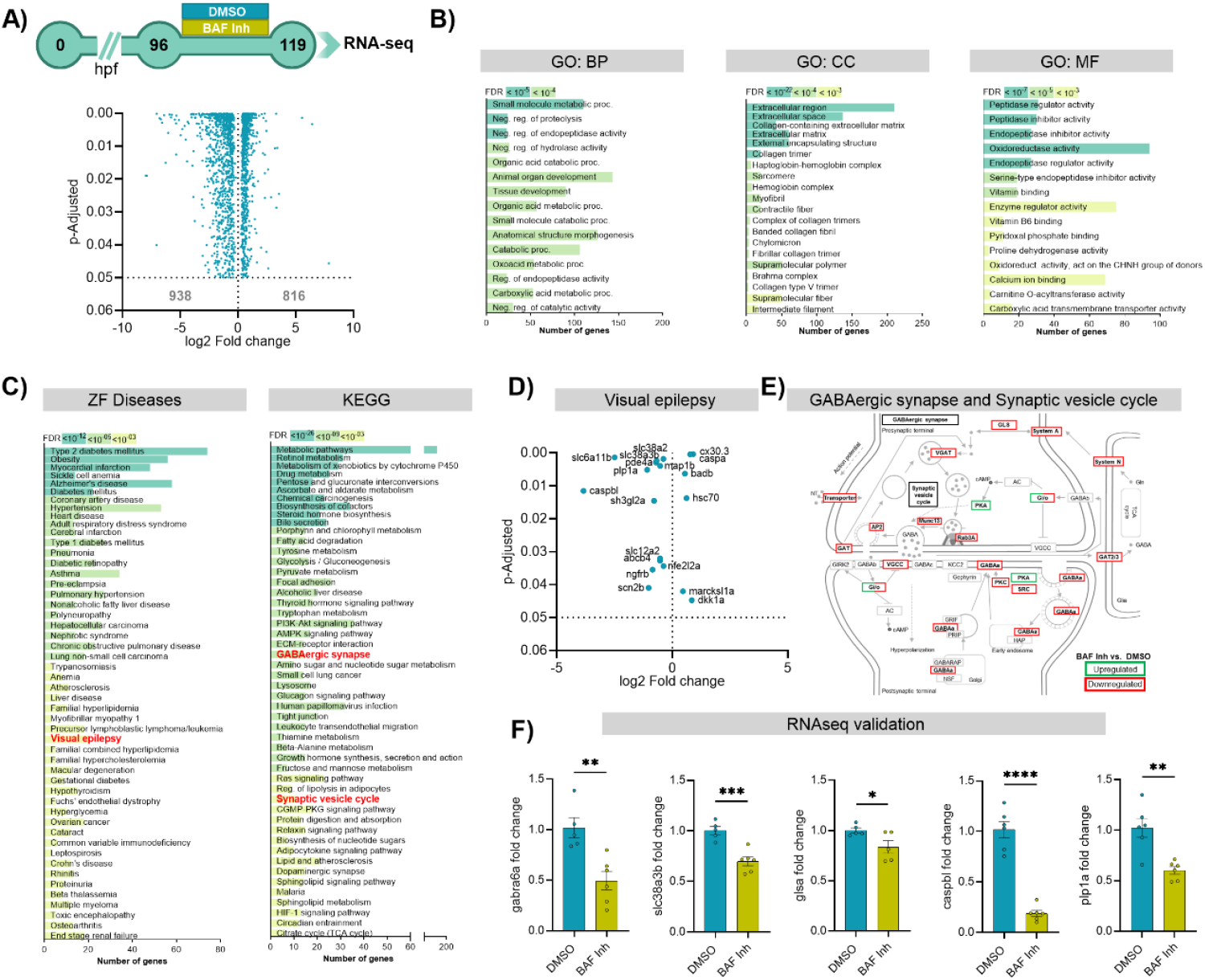
Transcriptome deregulation following BAF complex inhibition. **(A)** Experimental timeline. Zebrafish larvae were treated at 96 hours post-fertilization (hpf) with either DMSO or BAF complex inhibitor (BAF Inh, 10 µM). At 119 hpf, larvae were sacrificed and their heads collected for RNA sequencing (n = 5; each replicate consisted of 30 heads). A volcano plot illustrates 1,754 differentially expressed genes (DEGs) following BAF inhibition, with 938 downregulated and 816 upregulated. Only genes with an adjusted *p*-value < 0.05 are shown. **(B)** Gene Ontology (GO) analysis of DEGs (both up- and downregulated) in the categories: Biological Process (GO:BP), Cellular Component (GO:CC), and Molecular Function (GO:MF), using a false discovery rate (FDR) cutoff of 0.05. **(C)** Zebrafish disease and the Kyoto Encyclopedia of Genes and Genomes (KEGG) pathway analyses of DEGs (both up- and downregulated) following BAF Inh (FDR cutoff: 0.05). **(D)** Volcano plot showing DEGs associated with the “visual epilepsy” pathway. **(E)** Modified KEGG pathway diagrams for “GABAergic synapse” and “Synaptic vesicle cycle” highlighting DEGs following BAF Inh. Genes upregulated are shown in green; downregulated genes in red. **(F)** Validation of selected DEGs from the RNA sequencing analysis in independent samples (*gabra6a*, t(9) = 3.85, *p* = 0.0039; *slc38a3b*, t(9) = 5.09, *p* = 0.0007; *glsa*, t(8) = 2.40, *p* = 0.0431; *capbl*, t(10) = 9.53, *p* < 0.0001; *plp1a*, t(10) = 4.20, *p* = 0.0018) (n_DMSO_ = 5-6, n_BAFInh_ = 5-6). Data are presented as mean ± standard error of the mean (SEM). \**p* < 0.05, \*\**p* < 0.01, \*\*\**p* < 0.001, \*\*\*\**p* < 0.0001.

The Gene Ontology (GO) analysis using significantly DEGs (Benjamini–Hochberg adjusted *p*-value < 0.05) identified enrichment in pathways related to small-molecule metabolism, negative regulation of enzymatic activity, and developmental processes. In Molecular Function ontology (MF) term related to calcium ion binding was enriched (**Fig. 2B**). Disease and pathway enrichment analyses further implicated metabolic disorders and neurological conditions: ZFIN Disease terms highlighted “visual epilepsy”, while the Kyoto Encyclopedia of Genes and Genomes (KEGG) pathways included “Synaptic vesicle cycling” and “GABAergic synapse” (**Fig. 2C**). The set of epilepsy-related genes showed a significant deregulation of epilepsy modulators involved in GABAergic tone, neuronal inflammation, and apoptosis (**Fig. 2D**). Also, mapping all DEGs in the “GABAergic synapse” and “Synaptic vesicle cycle” pathways using the KEGG database revealed widespread downregulation of presynaptic GABA transporters, vesicular loading machinery, receptor subunits and associated signaling molecules (**Fig. 2E**). To validate these findings, we performed RT-qPCR on independent samples for five representative genes from the visual epilepsy and GABAergic synapse pathways (*gabra6a, slc38a3b, glsa, caspbl, plp1a)* and observed fold changes consistent with the RNAseq data, with all genes significantly downregulated in BAF inhibitor treated larvae (**Fig. 2F**). This validates the RNAseq results and the deregulation of GABAergic synapse-related and epilepsy-related genes by the BAF inhibitor. We also investigated whether these molecular effects were a consequence of seizure activity or a direct result of BAF complex inhibition by analyzing gene expression with and without AP5 treatment to block seizures (**SI Appendix Fig. S5A i**). The analysis showed no rescue in gene expression by AP5, suggesting that the effects result from BAF inhibition rather than seizure activity (**SI Appendix Fig. S5A ii**).

To assess whether BAF complex inhibition engages conserved epilepsy-related transcriptional signatures, our RNAseq data were compared with a publicly available mouse hippocampal microarray data from a kainic acid (KA)–induced mesial temporal lobe epilepsy model (GEO accession GSE88992) (48). In this study, C57BL/6J mice received unilateral stereotactic KA or saline injections into the dorsal hippocampus to induce status epilepticus and were sacrificed at 6, 12, or 24 hours post-injection (**SI Appendix Fig. S6A**). Comparative analysis of KA-injected mouse and BAF inhibitor-treated zebrafish larvae transcriptomes identified 236 shared DEGs at 6 hours, 710 at 12 hours, and 450 at 24 hours (**SI Appendix Fig. S6B i–iii**). At 6 hours, 60 genes were commonly upregulated and 48 downregulated; at 12 hours, 154 were upregulated and 172 downregulated; and at 24 hours, 99 were upregulated and 101 downregulated in the same direction across datasets (**SI Appendix Fig. S6C i–iii**). Notably, Brg1 was downregulated at 12 hours, prompting us to focus subsequent pathway analyses on this time point (**SI Appendix Fig. S6B ii**). GO and KEGG enrichment analyses revealed conserved dysregulation of neuronal and synaptic processes across species, including enrichment of GABAergic synapse and synaptic vesicle cycle pathways, indicating shared GABAergic synaptic impairment in both BAF perturbation and the KA-induced epilepsy model (**SI Appendix Fig. S6D i–iv**).

Together, these results demonstrate that BAF complex inhibition triggers pro-epileptic transcriptional changes characterized by disrupted GABAergic signaling and synaptic vesicle cycling, which is conserved across species and recapitulated in an epilepsy model.

### GABAergic deficits underlie BAF complex inhibition-induced seizure-like behavior

Next, we sought to investigate the molecular mechanisms underlying the inhibition of the BAF complex. Among the neurotransmitter pathways deregulated in the RNAseq analysis, the GABAergic synapse was enriched. We investigated whether the GABAergic system, compared to the glutamatergic system, was specifically impaired by BAF loss and thereby contributed to the altered balance. To this end, whole-mount immunostaining was performed for glutamatergic and GABAergic synaptic markers in larvae treated with the BAF inhibitor for 24 hours (**Fig. 3A i**). First, we analyzed the number of nuclei in the OT to determine whether 24 hours inhibition led to a reduction in cell number. Analysis showed no significant change in overall cell count, as indicated by nuclei staining (**Fig. 3A ii**). We then examined synaptic markers. No difference was observed in the intensity of the excitatory marker vGlut1/2 (**Fig. 3A iii**). However, we detected a significant decrease in the GABAergic neurons marker GAD65/67 and the postsynaptic inhibitory marker Gephyrin in BAF inhibitor-treated larvae (**Fig. 3A iv, v**), suggesting a reduction in inhibitory synaptic signaling.

**Figure 3.**
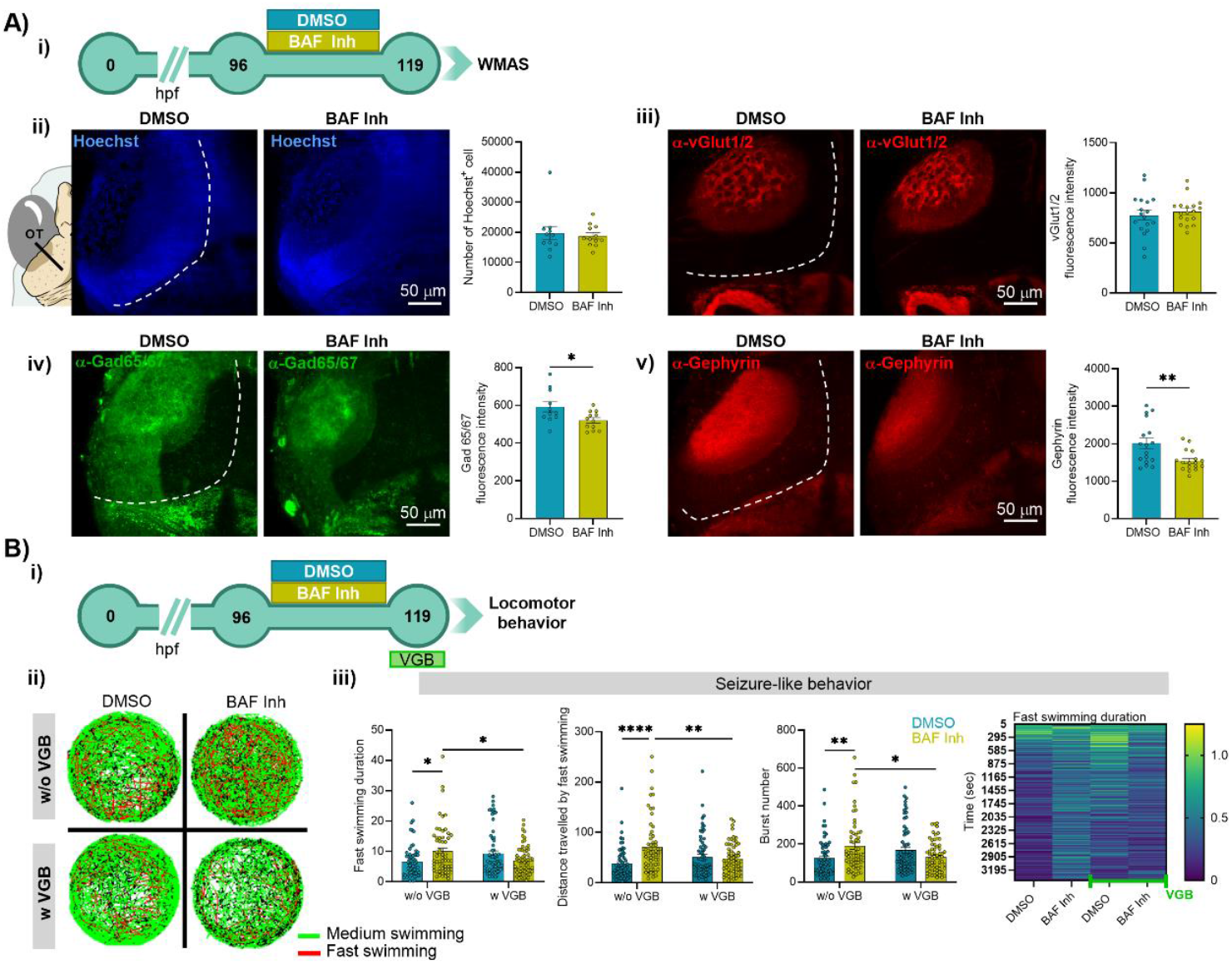
BAF complex inhibition affects the GABAergic system, and the seizure-like behavior is rescued by anti-seizure drug Vigabatrin. **(A) i)** Experimental timeline. Zebrafish larvae were treated at 96 hours post-fertilization (hpf) with either DMSO or a BAF complex inhibitor (BAF Inh, 10 µM) overnight and then were used for whole-mount immunostaining. **ii)** Representative Z-projection (maximum intensity) images showing nuclei in the optic tectum (OT) of larvae treated with DMSO or BAF Inh. Quantification of cell nuclei in the OT of DMSO and BAF Inh treated zebrafish (Mann-Whitney test, U = 63, *p* = 0.8801), (n_DMSO_ = 11, n_BAFInh_ = 12). Data are presented as mean ± standard error of the mean (SEM). **iii)** Representative Z-projection (maximum intensity) images showing vGlut1/2 staining in the OT of larvae treated with DMSO or BAF Inh. Quantification of vGlut1/2 in the OT of DMSO and BAF Inh treated zebrafish (t(34) = 0.64, *p* = 0.5267), (n_DMSO_ = 18, n_BAFInh_ = 18). Data are presented as mean ± (SEM). **iv)** Representative Z-projection (maximum intensity) images showing GAD65/67 level in the OT of larvae treated with DMSO or BAF Inh. Quantification of GAD65/67 in the OT of DMSO and BAF Inh treated zebrafish (t(21) = 2.45, *p* = 0.0232), (n_DMSO_ = 11, n_BAFInh_ = 12). Data are presented as mean ± SEM. **v)** Representative Z-projection (maximum intensity) images showing gephyrin staining in the OT of larvae treated with DMSO or BAF Inh. Quantification of gephyrin in the OT of DMSO and BAF Inh treated zebrafish (t(23.03) = 3.052, *p* = 0.0057, Welch’s correction), (n_DMSO_ = 17, n_BAFInh_ = 17). Data are presented as mean ± SEM. **(B) i)** Experimental timeline. Zebrafish larvae were treated at 96 hpf with either DMSO or BAF Inh overnight, followed by a 2-hour treatment with vigabatrin (VGB, 60 µM) before assessment of their locomotor behavior. **ii)** Representative locomotor tracks of larvae treated with DMSO or BAF Inh, with and without VGB. Medium swimming track (0.5-2 cm/sec) is shown in green, while fast swimming track (>2 cm/sec) is shown in red. **iii)** Quantification of seizure-like behavior parameters: Fast swimming duration (two-way ANOVA: Treatment vs. VGB interaction, F (1, 246) = 12.31, *p* = 0.0005; Tukey post hoc test), distance traveled by fast swimming (two-way ANOVA: Treatment vs. VGB interaction, F (1, 247) = 13.29, *p* = 0.0003, Treatment effect: F (1, 247) = 8.363, *p* = 0.0042; Tukey post hoc test), number of burst events (two-way ANOVA: Treatment vs. VGB interaction, F (1, 247) = 12.60, *p* = 0.0005; Tukey post hoc test) and a heatmap illustrating fast swimming duration over time (n_DMSO_ = 61, n_BAFInh_ = 61, n_DMSO_VGB_ = 65, n_BAFInh_VGB_ = 64). Data are presented as mean ± SEM. \**p* < 0.05, \*\**p* < 0.01, \*\*\*\**p* < 0.0001.

To confirm the GABA dependency, larval behavior after 24 hours of BAF inhibitor treatment followed by 2 hours of vigabatrin (VGB) exposure was assessed (**Fig. 3B i**). VGB is a GABA-transaminase inhibitor that prevents GABA degradation, commonly used as an anti-seizure drug in humans (49). As observed previously, BAF inhibited fish (without VGB) exhibited increased seizure-like activity (**Fig. 3B ii, iii**). Remarkably, VGB treatment abolished these increases, restoring all measures of seizure-like behavior to control levels (**Fig. 3B ii, iii**).

These results show that BAF inhibition specifically disrupts the GABAergic system, leading to the imbalance between excitatory and inhibitory signaling. This is further confirmed by the rescue of seizure-like behavior by VGB, underscoring its potential as a treatment for BAF-related seizures.

### The Brg1 subunit of the BAF complex regulates seizure susceptibility in zebrafish

To determine whether the effects induced by BAF complex inhibition were specifically due to the Brg1 subunit, we generated a zebrafish line with a targeted deletion of the *smarca4* gene (Brg1 in zebrafish) using the CRISPR/Cas9 system (**SI Appendix Fig. S7A–F**). Predicted mutation effects were confirmed by significantly reduced *smarca4* expression at mRNA and protein levels in heterozygote (HET) and knockout (KO) larvae (**SI Appendix Fig. S7H–I**). Phenotypically, KO showed smaller eyes, abnormal posture, and severe cardiac edema, while wild-type (WT) and HET appeared normal (**SI Appendix Fig. S7J**), consistent with previous Brg1 disruption studies (50, 51).

First, the STFBC behavior at 24 hpf in the *smarca4* mutant line was assessed (**Fig. 4A i, ii**). Quantification of total coils showed no significant difference between HET and WT fish, whereas a significant reduction in coiling was observed in KO embryos compared to both WT and HET (**Fig. 4A iii**). Similar to our findings with BAF inhibitor, HET embryos exhibited a significant increase in double coils compared to WT, while KO embryos showed fewer coils overall compared to both WT and HET. These results show that partial loss of the Brg1 subunit contributes to early hyperexcitability (**Fig. 4A iii**). To confirm this, we performed a rescue experiment by injecting mouse *Brg1* (m*Brg1*) RNA (52) at the one-cell stage and then examined STFBC behavior at 24 hpf. The m*Brg1* injection rescued the double coil impairment in HET and thereby confirmed the specificity of Brg1 in regulating the neuronal hyperexcitability at this developmental stage (**Fig. 4A iv**).

**Figure 4.**
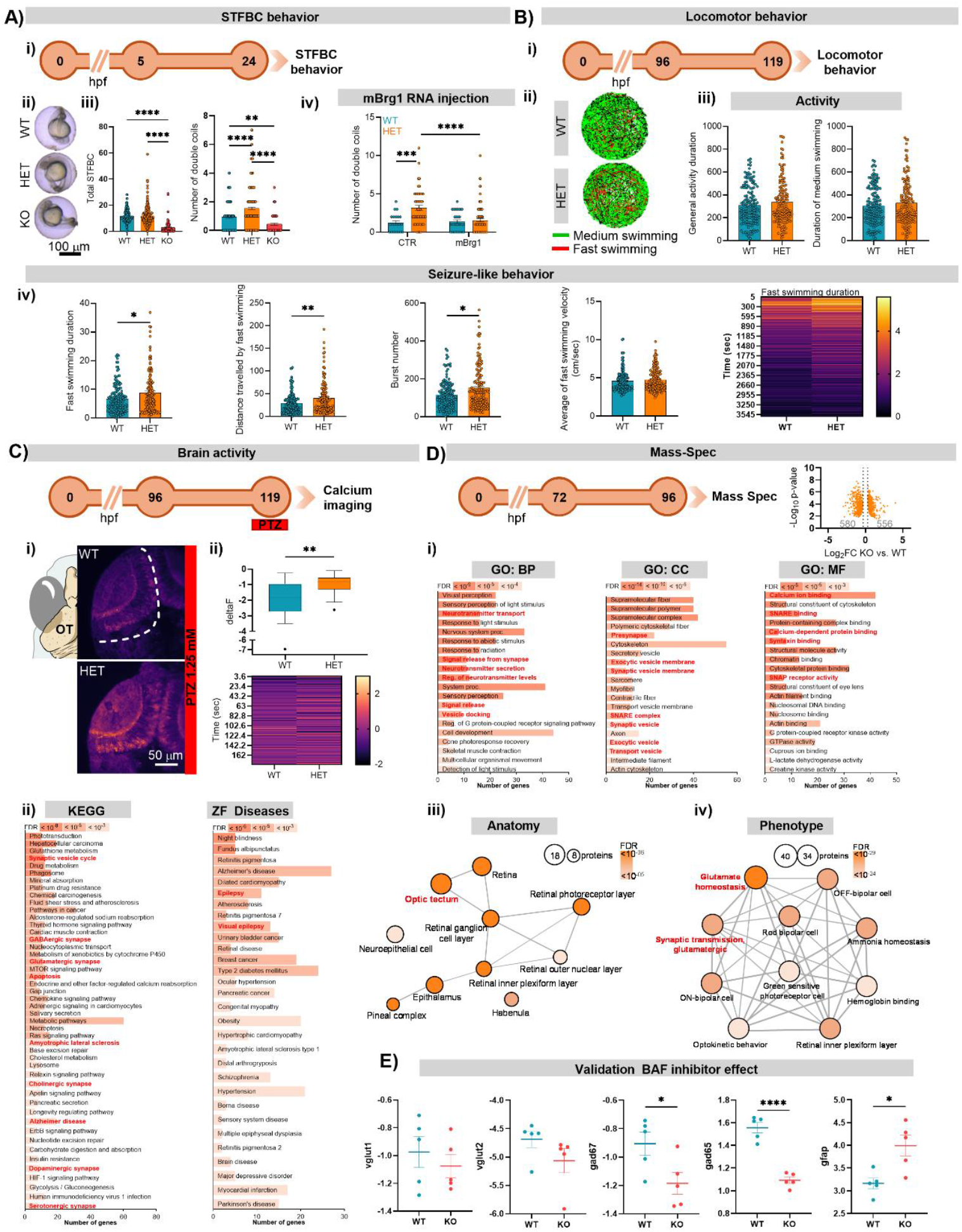
The CRISPR/Cas9-generated *smarca4*-specific mutant line recapitulates the effects observed with BAF inhibitor treatment. **(A) i)** Experimental timeline. Spontaneous tail flick and body coiling (STFBC) behavior in *smarca4* mutants was measured at 24 hours post-fertilization (hpf). **ii)** Representative images of *smarca4* mutant larvae at 24 hpf during the recording of STFBC behavior. **iii)** Quantification of total coils (Kruskal-Wallis test, H = 169.3, *p* < 0.0001; Dunn’s post hoc test) and double coils (Kruskal-Wallis test, H = 57.68, *p* < 0.0001; Dunn’s post hoc test) (n_WT_ = 224, n_HET_ = 311, n_KO_ = 91). Data are presented as mean ± standard error of the mean (SEM). **iv)** Quantification of total coils (Kruskal-Wallis test, H = 18.60, *p* < 0.0001; Dunn’s post hoc test) and double coils (Kruskal-Wallis test, H = 3.492, *p* = 0.1745; Dunn’s post hoc test) in the *smarca4* line injected with mouse Brg1 (m*Brg1*) RNA (n_WT_ = 30, n_HET_ = 87). Data are presented as mean ± SEM. **(B) i)** Experimental timeline. Locomotor behavior of *smarca4* mutant line was assessed at 119 hpf. **ii)** Representative locomotor tracks of *smarca4* mutant larvae. **iii)** General locomotor activity: Duration of general activity (Mann-Whitney test, U = 10234, *p* = 0.437) and duration of medium swimming (Mann-Whitney test, U = 10275, *p* = 0.472). Data are presented as mean ± SEM. **iv)** Quantification of seizure-like behavior parameters: Fast swimming duration (Mann-Whitney test, U = 9095, *p* = 0.0193), distance traveled by fast swimming (Mann-Whitney test, U = 8779, *p* = 0.0055), number of burst events (Mann-Whitney test, U = 9040, *p* = 0.0158), average velocity during fast swimming (Mann-Whitney test, U = 9837, *p* = 0.2191) and a heatmap illustrating fast swimming duration over time (n_WT_ = 149, n_HET_ = 144). Data are presented as mean ± SEM. **(C)** Experimental timeline. Calcium imaging was performed following a 3-minute incubation with a subthreshold dose of pentylenetetrazole (PTZ, 1.25 mM) on *smarca4* mutant larvae. **i)** Representative Z-projection (maximum intensity) images showing neuronal activity in the OT of *smarca4* larvae. **ii)** Quantification of neuronal activity (Mann-Whitney test, U = 105.5, *p* = 0.0011) and heatmap showing temporal dynamics of neuronal activity (n_WT_PTZ_ = 20, n_HET_PTZ_ = 24). Data are presented as Tukey box-and-whisker plots. **(D)** Experimental timeline. Mass spectrometry was conducted on *smarca4* mutant line at 96 hpf (n_WT_ = 5, n_HET_ = 5, each replicate consisted of 5 heads). A volcano plot illustrates 1136 deregulated proteins in *smarca4* mutant line, with 580 downregulated and 556 upregulated. Only significantly deregulated proteins (Student’s t-test, 2-sided, permutation-based FDR = 0.001, S0 = 0.1) are shown. **i)** Gene Ontology (GO) analysis of deregulated proteins (both up- and downregulated) in the categories: Biological (GO:BP), Cellular Component (GO:CC), and Molecular Function (GO:MF), using a false discovery rate (FDR) cutoff of 0.05. **ii)** The Kyoto Encyclopedia of Genes and Genomes (KEGG) and zebrafish disease pathway analyses of deregulated proteins (both up- and downregulated) in *smarca4* mutant line (FDR cutoff: 0.05). **iii-iv)** Anatomy and phenotype databases analyses of deregulated proteins (both up- and downregulated) in the *smarca4* mutant line (the color intensity of the bubbles represents the FDR, while the size of the bubbles represents the number of proteins in each pathway). **(E)** Validation of the effect of BAF Inh on the glutamatergic system (protein abundance: vglut1: t(8) = 0.727, *p* = 0.4880; vglut2: Mann-Whitney test, U = 4, *p* = 0.095), the GABAergic system (protein abundance: gad67: t(8) = 2.483, *p* = 0.0379; gad65: t(8) = 8.546, *p* < 0.0001) and glial cells (protein abundance: GFAP: t(8) = 3.243, *p* = 0.0118) in the *smarca4* mutant line. Data are presented as mean ± SEM. \**p* < 0.05, \*\**p* < 0.01, \*\*\**p* < 0.001, \*\*\*\**p* < 0.0001.

We next evaluated locomotor behavior at 119 hpf in WT and HET larvae. KO larvae were not included at this stage due to lethality associated with *smarca4* KO (**Fig. 4B i**). Locomotor tracking showed no differences in basal locomotion between HET and WT larvae (**Fig. 4B ii, iii**). However, HET larvae displayed elevated seizure-like behaviors (**Fig. 4B iv**).

To further validate the specific role of Brg1 in neuronal excitability, we performed calcium imaging following administration of a subthreshold dose of PTZ in *smarca4* mutant line crossed with *Tg(HuC:GCaMP5G)* (**Fig. 4C**). HET larvae exhibited a significantly increased calcium response in the OT (**Fig. 4C i, ii**), indicating heightened neuronal activity.

To investigate the molecular changes associated with the observed phenotypes, mass spectrometry-based proteomic profiling was performed at 96 hpf in WT and KO *smarca4* zebrafish (**Fig. 4D**). Differential expression analysis revealed a substantial number of deregulated proteins in KO embryos compared to WT, with 580 proteins downregulated and 556 upregulated (**Fig. 4D**). The GO enrichment analysis using significantly deregulated proteins (Student’s t-test, 2-sided, permutation-based FDR = 0.001, S0 = 0.1) revealed strong enrichment in pathways related to synaptic transmission, ion transport, and signal transduction. Notably, proteins involved in glutamate signaling, calcium ion transport, and neurotransmitter release were significantly altered (**Fig. 4D i**).

Pathway enrichment analysis using KEGG and Zebrafish Disease databases further confirmed disruptions in key neurological and signaling pathways. These included GABAergic and glutamatergic synapses, calcium signaling, and apoptosis, as well as associations with neurological disorders such as epilepsy and visual epilepsy (**Fig. 4D ii**).

Anatomical mapping of the differentially expressed proteins showed enrichment in brain regions involved in visual processing and synaptic integration, including the retina and habenula. Interestingly, the largest protein cluster was associated with the OT, suggesting this structure is particularly affected by *smarca4* loss and consistent with BAF inhibitor results (**Fig. 4D iii**). Phenotypic annotation highlighted disruptions in glutamate homeostasis and synaptic transmission pathways (**Fig. 4D iv**). Finally, protein analyses further supported the immunostaining results obtained from the BAF inhibitor experiments. Compared with WT, KO larvae exhibited reduced levels of GABAergic proteins (GAD65, GAD67), unchanged glutamatergic proteins (vGlut1, vGlut2), and elevated GFAP expression (**Fig. 4E**).

Together, these results demonstrate that loss of Brg1 specifically induces early-onset neural hyperexcitability and seizure-like behaviors, accompanied by widespread molecular and cellular alterations, including GABAergic dysfunction. This underscores Brg1’s crucial role in neural development and in maintaining the excitatory/inhibitory balance underlying seizure susceptibility.

### Treatment with the active form of Vitamin B6 reduces seizure-like behavior in *smarca4* mutants

To investigate potential molecular mechanisms underlying the seizure-like behavior observed in the s*marca4* line, we performed GO MF enrichment analysis using less stringent parameters than the previous analysis. Specifically, all proteins with a *q*-value less than 0.05 were used as input for the analysis.

The GO enrichment analysis revealed significant clustering related to the actin cytoskeleton, chromatin binding, calcium binding, and various metabolic processes. Notably, in *smarca4* KO, a distinct functional module associated with vitamin binding was identified, encompassing the GO terms “Vitamin binding”, “Vitamin B6 binding”, and “Pyridoxal phosphate binding” (**Fig. 5A**). A similar enrichment pattern was observed in the RNAseq data following BAF inhibitor treatment (GO MF, **Fig. 2B**).

**Figure 5.**
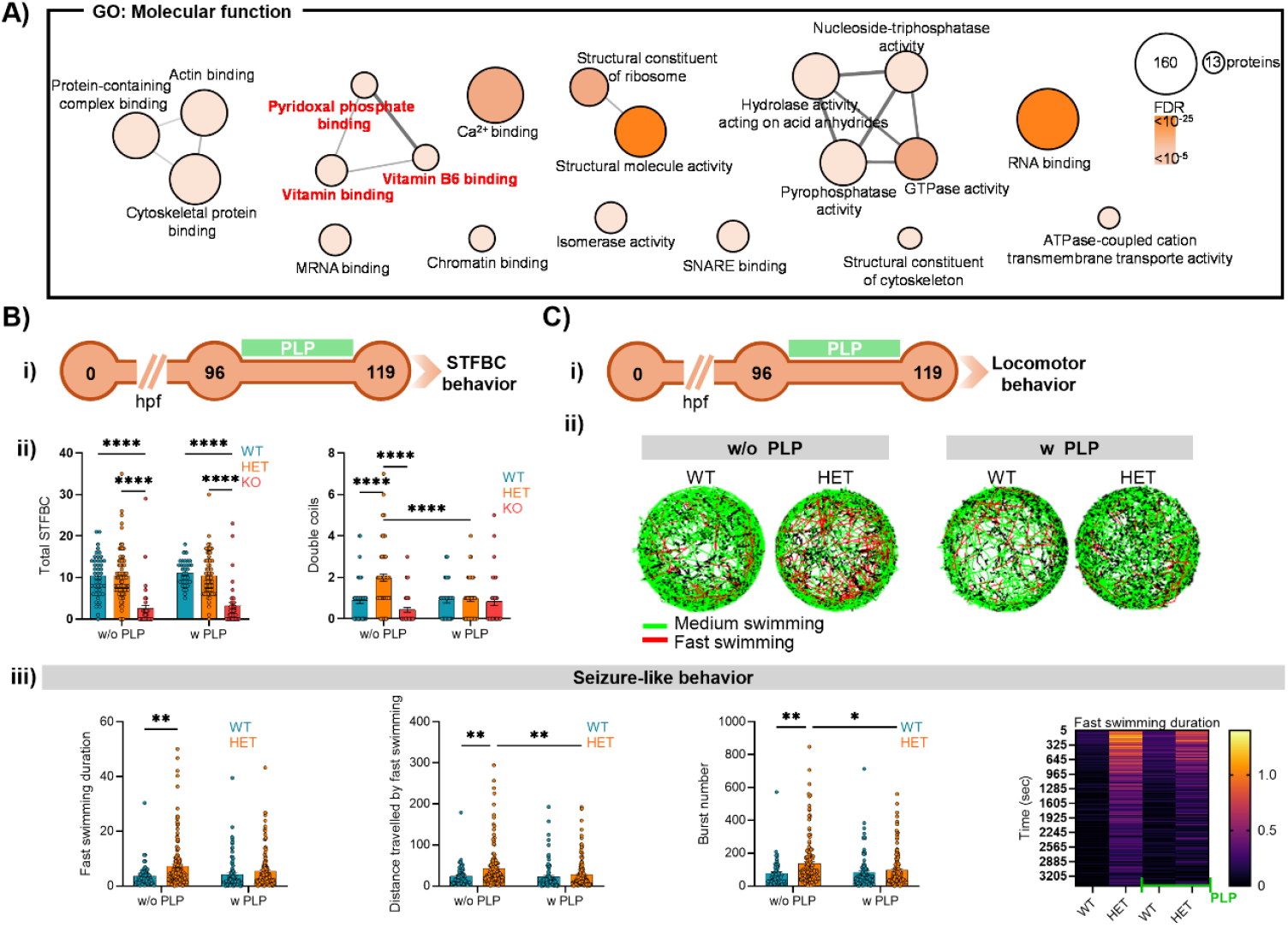
Supplementation with the active form of vitamin B6, pyridoxal 5’-phosphate (PLP), rescues the seizure-like behavior in the *smarca4* mutant line. **(A)** Gene Ontology (GO) analysis of deregulated proteins (both up- and downregulated, *q*-value < 0.05) in the category: Molecular Function (GO:MF), using a false discovery rate (FDR) cutoff of 0.05. **(B) i)** Experimental timeline. Pyridoxal phosphate (PLP) supplementation (100 µM) was administered at 5 hours post-fertilization (hpf) to *smarca4* mutant embryos, and spontaneous tail flick and body coil (STFBC) behavior was assessed at 24 hpf. **ii)** Quantification of total coils (two-way ANOVA: Genotype effect, F (2, 307) = 70.17, *p* < 0.0001, Tukey post hoc test) and double coils (two-way ANOVA: Genotype effect, F (2, 307) = 16.16, *p* < 0.0001; Genotype vs. Treatment interaction, F (2, 307) = 11.61, *p* < 0.0001; Tukey post hoc test) in *smarca4* mutant line (n_WT_ = 40, n_HET_ = 79, n_KO_ = 46, n_WT_PLP_ = 34, n_HET_PLP_ = 68, n_KO_PLP_ = 41). Data are presented as mean ± standard error of the mean (SEM). **(C) i)** *smarca4* mutant larvae were treated with PLP at 96 hpf and incubated overnight. Locomotor behavior was subsequently assessed. **ii)** Representative locomotor tracks of *smarca4* mutant larvae with and without PLP treatment. Medium swimming track (0.5-2 cm/sec) is shown in green, while fast swimming track (>2 cm/sec) is shown in red. **iii)** Seizure-like behavior parameters: Fast swimming duration (two-way ANOVA: Genotype effect: F (1, 473) = 11.42, *p* = 0.0008; Tukey post hoc test), distance traveled with fast swimming (two-way ANOVA: PLP effect: F (1, 473) = 4.573, *p* = 0.033; Genotype effect: F (1, 473) = 9.648, *p* = 0.0020; Tukey post hoc test), number of burst episodes (two-way ANOVA: PLP vs. Genotype interaction: F (1, 473) = 4.167, *p* = 0.0418; Genotype effect: F (1, 473) = 9.900, *p* = 0.0018; Tukey post hoc test) and a heatmap illustrating fast swimming duration over time (n_WT_ = 62, n_HET_ = 163, n_WT_PLP_ = 90, n_HET_PLP_ = 162). Data are presented as mean ± SEM. \**p* < 0.05, \*\**p* < 0.01, \*\*\*\**p* < 0.0001.

Since vitamin B6 is known to function as a cofactor in neuronal transmission, particularly in GABAergic synapses (53–55), we investigated whether supplementation with pyridoxal phosphate (PLP, the active form of vitamin B6) could ameliorate the impairments observed in the *smarca4* mutant line.

STFBC behavior was assessed 24 hours after PLP treatment (**Fig. 5B i**). Quantification showed that total coil events were not affected in *smarca4* HET and KO larvae following PLP treatment (**Fig. 5B ii**). However, the number of double coils, which was elevated in HET larvae, was significantly reduced after PLP treatment to the same level as control (**Fig. 5B ii**).

Locomotor activity and seizure-like behavior were also assessed in 119 hpf larvae after 24 hours of PLP exposure (**Fig. 5C i**). Analysis of seizure-like behavior demonstrated that PLP treatment significantly reduced the characteristic increase in fast swimming observed in HET larvae compared to WT (**Fig. 5C ii, iii**).

These effects were also confirmed in an independent experiment using a specific pharmacological inhibitor of Brg1, IV255. Zebrafish treated with IV255 exhibited increased seizure-like behavior, similar to *smarca4* HET and zebrafish treated with the BAF inhibitor (**SI Appendix Fig. S8A i–iv**). Furthermore, co-treatment with PLP rescued this phenotype (**SI Appendix Fig. S8B i–iii**).

These findings demonstrate that the behavioral deficits caused by Brg1 reduction are responsive to PLP supplementation. This highlights the potential of targeting vitamin B6 as a therapeutic strategy for neurological dysfunctions associated with BAF complex activity reduction.

## Discussion

Epilepsy is increasingly viewed as a disease of disturbed network balance, shaped by changes in gene regulation, including epigenetic mechanisms (5, 6). In this study, we identify the chromatin remodeler Brg1 (Smarca4), the catalytic subunit of BAF complexes, as an important regulator of inhibitory neurotransmission and seizure susceptibility. Although mutations in BAF components are strongly associated with neurodevelopmental disorders that often include epilepsy, the mechanisms linking chromatin remodeling to seizures have remained unclear (7, 13, 18, 56, 57). Using pharmacological inhibition and genetic disruption of *smarca4* in zebrafish, we show that partial loss of Brg1 function is sufficient to induce neuronal hyperexcitability and seizure-like behavior.

A key finding of our work is the selective impairment of the GABAergic system following Brg1 reduction. Our transcriptomic and proteomic analyses both revealed deregulation of pathways involved in GABAergic synapses and synaptic vesicle cycling. At the same time, markers of glutamatergic transmission remained essentially unchanged. This pattern fits with previous studies showing that Brg1 and BAF complexes regulate specific neuronal genes rather than acting as global transcriptional regulators (11, 14, 29). Developmental RNA sequencing studies also reported changes in GABA-related pathways after Brg1 loss (58), and Brg1 deletion in postnatal neurons was shown to impair synapse development and function (28). Moreover, studies on the ultraconserved transcription-regulating long non-coding RNA (lncRNA) Evf2 demonstrated that Evf2 directly interacts with and inhibits Brg1 chromatin-remodeling activity at Dlx enhancers critical for GABAergic interneuron development (59). Notably, mutations associated with Coffin–Siris syndrome localize to the Brg1 RNA-binding and DLX1-binding domains (21, 59, 60), supporting the idea that disruption of RNA-dependent Brg1 regulation may be particularly detrimental for inhibitory circuit formation and, as suggested by our findings, could contribute to the neuronal hyperexcitability and seizure-like behavior. Importantly, the seizure-like behavior observed in our model was rescued by vigabatrin, which increases brain GABA levels by inhibiting GABA transaminase (61, 62). This functional rescue supports the conclusion that reduced inhibitory tone is a major driver of hyperexcitability following Brg1 disruption.

Our data further indicate that Brg1-related seizure susceptibility is sensitive to the amount of loss of functional protein. Full knockout of *smarca4* caused severe developmental defects and early lethality. On the other hand, heterozygous mutants displayed robust seizure-like phenotypes without significant baseline locomotor abnormalities. This closely reflects the clinical situation, where many patients with Coffin–Siris syndrome or related conditions carry heterozygous variants in SWI/SNF genes and frequently present with epilepsy (20–24, 60, 63). Our results, therefore, provide functional support for the idea that partial impairment of Brg1-dependent chromatin remodeling can be sufficient to destabilize neuronal network activity.

At the circuit level, Brg1 reduction increased neuronal activity in the optic tectum, as shown by calcium imaging and electrophysiological recordings. This observation is consistent with earlier work demonstrating that Brg1 regulates activity-dependent gene expression, including the control of immediate-early genes such as *c-fos* (14). In zebrafish, seizures often initiate in midbrain regions, and the optic tectum is particularly sensitive to epileptiform activity (64, 65). Therefore, the brain region examined in this study aligns well with those used in established zebrafish epilepsy models. Prolonged inhibition of the BAF complex led to neuroinflammation, glial activation, and neuronal loss. Similar phenotypes have been reported after Brg1 deletion in mammalian systems (58, 66). In our experiments, these changes appeared after the onset of hyperexcitability, suggesting that inflammation and degeneration are likely downstream consequences of sustained network dysfunction rather than primary causes of seizure initiation. This interpretation is also consistent with transcriptomic signatures observed in human epilepsies, including mesial temporal lobe epilepsy (42, 43).

Finally, unbiased omics analyses pointed to altered expression of proteins involved in vitamin B6 and pyridoxal 5′-phosphate (PLP) binding. Vitamin B6 is a known cofactor for enzymes involved in neurotransmitter metabolism and has long been linked to epilepsy, particularly through effects on GABA synthesis and turnover (53, 67–72). Supplementation with PLP rescued seizure-like behavior in both genetic and pharmacological Brg1 loss models. Although PLP is not specific to Brg1-dependent pathways, its rescue effect is consistent with impaired GABAergic transmission being a central feature of the phenotype. Experimental studies have shown that vitamin B6 availability can directly influence GABA synthesis and degradation (73), supporting this interpretation.

In summary, our findings identify Brg1 as a regulator of inhibitory neurotransmission and seizure susceptibility. Moreover, by linking Brg1 to control of GABAergic transmission, this study provides a mechanistic explanation for the frequent association between mutations in BAF complex subunits and epilepsy.

## Materials and Methods

Detailed materials and methods are in the **Supplementary Methods** in the **SI Appendix**.

### Zebrafish breeding

Zebrafish used in this study, with the exception of non-invasive local field potential recording, were at developmental stages below the equivalent of 120 hours post-fertilization (hpf) and of unspecified sex. The non-invasive local field potential recordings were conducted at developmental stages ≥120 hpf, and the experimental protocol was approved by the Ethics Committee of KU Leuven (approval number P034/2024) and by the Belgian Federal Department of Public Health, Food Safety, and Environment (approval number LA1210261). Zebrafish were bred according to international standards at the International Institute of Molecular and Cell Biology, Warsaw, Poland or KU Leuven, Leuven, Belgium. All experimental procedures were carried out in accordance with the Act of 15 January 2015 on the protection of animals used for scientific and educational purposes and with Directive 2010/63/EU of the European Parliament and of the Council of 22 September 2010 on the protection of animals used for scientific purposes. Experiments were performed on wild-type (AB×TL strain), *Tg(HuC:GCaMP5G*) (74), *Tg(mpeg1:mCherry*) (75) and *smarca4* lines.

### *smarca4* mutant line generation

The *smarca4* mutant line was generated using the CRISPR/Cas9 system. A 20-base pair (bp) sequence (gRNA: 5’-GGTGCTATGCCTATGGAAGG-3’) was designed to target exon 3 of the *smarca4* coding region. Cas9 mRNA was synthesized from the pT3TS-nlsCas9nls plasmid (Addgene, #46757) (76). A mixture of gRNA and Cas9 mRNA was injected into one-cell stage zebrafish embryos.

### RNA sequencing

Paired-end sequencing (2 × 150 bp) was performed on Illumina NovaSeq 6000 using NovaSeq 6000 S4 flow cell (Illumina) to target a depth of 25–30 million reads per sample at the Laboratory of Sequencing of the Nencki Institute, Warsaw, Poland.

### Behavioral analysis

Spontaneous tail flick and body coil behavior was measured by counting the total and double coils in 24 hpf larvae (37, 38, 77, 78). For locomotor behavior, larvae were recorded in the Zebrabox system (ViewPoint) and recorded for 1 hour in darkness (37).

### Brain activity analysis

Calcium imaging was performed using a Zeiss Lightsheet microscope (40X) to acquire 3D time-lapse recordings of the optic tectum in Tg(*HuC:GCaMP5G*) larvae. For non-invasive local field potential (LFP) recordings, electrical activity was recorded from the optic tectum and analyzed by Welch’s method for power spectral density (PSD) (42, 79).

### Mass spectrometry

Snap-frozen heads from 96 hpf *smarca4* mutant larvae were submitted for protein identification using liquid chromatography–tandem mass spectrometry (LC–MS/MS). Measurements were performed at the Proteomics Core Facility, IMol Polish Academy of Sciences, Warsaw, Poland.

### Statistical analysis

All statistical analyses were performed using GraphPad Prism 10 software. Exact sample sizes for each experiment are indicated in the figure legends. Data are presented as mean ± standard error of the mean (SEM). A *p*-value of less than 0.05 was considered statistically significant. Tests and details of analysis can be found in the **Supplementary Methods** in the **SI Appendix**.

## Acknowledgments

We thank Dr. Majewski (from Jacek Kuźnicki lab, IIMCB) and Dr. M Orger for sharing the zebrafish *mpeg:mcherry* and *Tg[HuC:GCaMP5G]* lines, respectively, and the IIMCB Zebrafish Core Facility for assistance with breeding and setting up the fish. We are also grateful to Alina Zielinska and Marek Sarnacki from our laboratory for technical assistance and support, and Angelika Jocek and Katarzyna Orzol for laboratory management logistics. This research was financed under the NCN MAESTRO grant 2020/38/A/NZ3/00447 project. This research was also possible thanks to the IIMCB IN-MOL-CELL Infrastructure funded by the European Union – NextGenerationEU under National Recovery and Resilience Plan. IN-MOL-CELL Infrastructure was also funded by the European Union under Horizon Europe (Project 101059801 - RACE) and by RACE-PRIME project carried out within the IRAP programme of the Foundation for Polish Science co-financed by the European Union under the European Funds for Smart Economy 2021-2027 (FENG). Equipment for the proteomic LC-MS/MS measurements was funded by the ‘Regenerative Mechanisms for Health’ project MAB/2017/2 within the International Research Agendas program of the Foundation for Polish Science, co-financed by the European Union under the European Regional Development Fund.

