## Supplementary material for "Loss of Brg1 promotes seizure development via GABAergic system disruption": SI Appendix

### Supporting Information Text

#### Supplementary Methods

**Zebrafish breeding.** Experiments with the BAF inhibitor and Brg1-specific inhibitor (IV255) were performed on wild-type (AB × TL strain) larvae, calcium imaging on *Tg(HuC:GCaMP5G)* (kind gift from Dr. Michael Orger (1)) and microglia analysis on *Tg(mpeg1:mCherry)* fish (kind gift from Dr. Lukasz Majewski (2)). To independently assess Brg1 specificity using a genetic approach, experiments were also conducted on *smarca4* mutant line and *Tg(HuC:GCaMP5G)*x*smarca4* larvae. Zebrafish used in this study were at developmental stages below the equivalent of 120 hours post-fertilization (hpf) at 28 °C. Zebrafish larvae used for non-invasive local field potential recordings, performed at the KU Leuven, were notified to the Ethics Committee of KU Leuven for early developmental stages <120 hpf and approved by the Ethics Committee of KU Leuven (approval number P034/2024) and by the Belgian Federal Department of Public Health, Food Safety, and Environment (approval number LA1210261) for developmental stages ≥120 hpf. Offspring from at least two different parental pairs were used for each experiment. Zebrafish were bred according to international standards. All experimental procedures were carried out in accordance with the Act of 15 January 2015 on the protection of animals used for scientific and educational purposes and with Directive 2010/63/EU of the European Parliament and of the Council of 22 September 2010 on the protection of animals used for scientific purposes.

**Drugs treatment.** Brg1/BRM inhibitor (BAF inhibitor; #HY-119374), IV255 (HY-155888), and Pyridoxal 5'-phosphate monohydrate (#HY-W011727A) were purchased from MedChemExpress (Monmouth Junction, NJ, USA). Pentylenetetrazol (PTZ) and Vigabatrin (VGB) were obtained from Sigma-Aldrich, and D-2-amino-5-phosphonovalerate (AP5, #0106) was purchased from Tocris. All compounds were dissolved in DMSO or water according to the manufacturers' instructions. Compounds were appropriately diluted in fish water (E3) and administered to equal numbers of zebrafish larvae. DMSO (as the control for the BAF inhibitor) was used at a final concentration of 0.02%, and the BAF inhibitor at 10 µM. DMSO (as the control for IV255) was used at 0.2%, and IV255 at 10 µM. AP5 and PLP were used at 100 µM, while VGB was used at 120 µM. The controls for both conditions were treated with 0.02% DMSO along with the corresponding concentrations of AP5 and VGB. Fish were anesthetized with a 0.168 mg/mL solution of tricaine methanesulfonate. For euthanasia, an overdose of tricaine was used.

***smarca4* mutant line generation.** CRISPR/Cas9 system was used to generate the *smarca4* mutant line. The 20-base pairs (bp) sequence (gRNA: 5'-GGTGCTATGCCTATGGAAGG-3') directly upstream of 5′-NGG sequences (PAM sequences) in the coding region of *smarca4* gene (exon 3) was identified by E-CRISP design tool (www.e-crisp.org/E-CRISP). The sequence with the lowest probability of off-targets and high on-target efficiency was chosen and validated with another gRNA design tool from IDT (www.eu.idtdna.com).
For oligos synthesis, the gRNA was inserted into a forward oligo with the following structure: 5′-GAAATTAATACGACTCACTATAGGG(NN…NN)GTTTTAGAGCTAGAAATAGC-3′, where the underlined segment is the T7 promoter, *NN…NN* is the gRNA sequence (excluding PAM), and the trailing sequence overlaps with the tracrRNA. A universal reverse oligo was used: 5′-AAAAGCACCGACTCGGTGCCACTTTTTCAAGTTGATAACGGACTAGCCTTATTTTAACTTGCT ATTTCTAGCTCTAAAAC-3′. Forward and reverse oligos were annealed and extended using Q5 High-Fidelity DNA Polymerase (NEB, #M0491). Products were verified on a 2% agarose gel, purified by Wizard SV Gel and PCR clean-up system (Promega, #A9281), and subsequently used as templates for in vitro transcription with the Hiscribe Quick T7 High Yield RNA Synthesis kit (NEB, #E2050) following manufacturer instructions. After transcription, samples were treated with DNase I, cleaned up using the Monarch Spin RNA Cleanup Kit (NEB, #T2040), quantified via Nanodrop, and stored at -80 °C. Cas9 mRNA was synthesized by linearizing the pT3TS-nlsCas9nls plasmid (gift from Wenbiao Chen, Addgene, #46757; http://n2t.net/addgene:46757; RRID:Addgene_46757) (3), followed by gel purification and transcription using the mMESSAGE mMACHINE T3 Transcription Kit (Thermo Fisher, #AM1348). After DNase treatment, the mRNA was purified by Monarch Spin RNA Cleanup Kit, quantified, and stored at -80 °C. For CRISPR injections, gRNA (60 ng/µL) and Cas9 mRNA (500 ng/µL) were mixed in a total volume of 4 µL. One nanoliter of this mix was injected into the cell of one-cell stage zebrafish embryos from at least two clutches.

Embryos were screened for editing efficiency via high resolution melting (HRM; primers: PF: 5’-CATGATGCCCTCTGGTCCT-3’, PR: 5’-GCTCAACAGAACCGGAGTG-3’) and Sanger sequencing (Genomed, https://www.genomed.pl/; primers for amplification and sequencing PF**:** 5’-CTCTGGGTGGTTCGGATCAC-3’, PR: 5’-CCATGATCTGTGCCCGCA-3’) after 24 h, following genomic DNA extraction. Once the founder fish was identified, it was crossed with wild-type zebrafish (AB × TL strain) to generate a stable line.

**RNA sequencing.** To obtain high-quality total RNA, 30 larval heads were pooled together and sonicated in the DNA/RNA protection reagent from the Monarch Total RNA Miniprep Kit (NEB, #T2110). Total RNA was extracted following the manufacturer's protocol. The RNA integrity was assessed by the Tapestation 2200 together with respective High Sensitivity RNA ScreenTape assay (Agilent Technologies). The average RNA Integrity Number equivalent (RINe) for samples used for downstream analysis was 7.7 or above.

In order to synthesize sequencing libraries, a two-step approach was applied. Total RNA was subjected to mRNA enrichment using NEBNext Poly(A) mRNA Magnetic Isolation Module (#E7490, NEB) followed by NEBNext Ultra II Directional RNA Library Prep Kit for Illumina (#E7760, NEB) according to the manufacturer's guidelines with minor adjustments. In brief, 1 µg of total RNA was used to enrich for polyadenylated RNA fraction. The obtained RNA was fragmented for 12 minutes at 94 °C to target insert size around 400 bp followed by extended first and second strand synthesis. To enrich for adaptor-ligated molecules, samples were pre-amplified with 4 cycles and left aside on ice. In order to determine an optimal number of PCR cycles and limit the possibility of generating PCR duplicates and artifacts, a subsequent qPCR reaction on previously pre-amplified samples was performed. Final libraries were validated in terms of quality and quantity by Quantus fluorometer (Promega) and Tapestation 2020 High Sensitivity D1000 assay (Agilent Technologies), respectively. Paired-end sequencing (2 x 150 bp) was performed on Illumina NovaSeq 6000 using NovaSeq 6000 S4 flow cell (Illumina) to target a depth of 25–30 million reads per sample.

Raw sequencing data were first quality-checked using FastQC (v. 0.11.9) (4) and then processed with Trimmomatic (v. 0.39) (5) to remove adapter contamination and low-quality reads. The cleaned reads were mapped to the Genome Reference Consortium Zebrafish Build 11 (GRCz11) using the Salmon algorithm (v. 0.14.1) (6). Quality control metrics included per-sequence quality score distributions, mean per-base quality scores, and sequence duplication levels. All samples showed consistently high-quality scores across read positions, with minimal variation between samples. Principal component analysis (PCA) was performed to evaluate sample clustering by condition (**SI Appendix Fig. S4**). Data normalization and statistical analysis were performed using the DESeq R package (v. 1.32.0). Significantly deregulated genes (Benjamini–Hochberg adjusted $p$-value < 0.05) identified from the RNA sequencing analysis were used as input for pathway analysis. Functional enrichment analysis was conducted using ShinyGO (v 0.76) (7) to identify affected pathways across Gene Ontology categories—Biological Processes (GO:BP), Cellular Component (GO:CC), and Molecular Function (GO:MF)—as well as in the Zebrafish disease database and the Kyoto Encyclopedia of Genes and Genomes (KEGG). For the KEGG analysis, gene IDs were converted from zebrafish to human to improve annotation quality and reveal additional potential pathways.

**Microarray reanalysis of a kainic acid-induced mesial temporal lobe epilepsy mouse model with comparison to BAF inhibitor-induced transcriptomic changes.**

In the study by Kalozoumi et al. (2018) (8), C57BL/6J mice that developed *status epilepticus* (SE) following dorsal hippocampal kainic acid (KA) injection were used to generate microarray-based transcriptomic data (data accessible at NCBI GEO database (9, 10), accession GSE88992). Mice were sacrificed at three time points: 6, 12, or 24 hours post-injection.

Raw Affymetrix Mouse Genome 430 2.0 CEL files corresponding to KA- and saline-injected samples were reprocessed in RStudio (v2021.09.0+351; https://posit.co/). Samples were grouped by experimental condition and analyzed separately for each post-injection time point (6 h, 12 h, 24 h). Raw intensity files were processed using robust multi-array average (RMA) normalization with the *oligo* package. Differential expression between KA and saline conditions was assessed using *limma* with empirical Bayes-moderated linear models. Resulting $p$-values were corrected for multiple testing using the Benjamini-Hochberg false discovery rate (FDR), and probes with FDR <

0.05 were significant. Probe sets were annotated to mouse gene symbols using the mouse4302.db Bioconductor package.

For cross-species comparison, zebrafish treated with BAF inhibitor differential expression dataset was used. Zebrafish genes were mapped to mouse orthologs based on Ensembl gene identifiers. Ortholog-mapped zebrafish DEGs were compared with mouse DEGs to assess conserved transcriptional responses. Shared differentially expressed genes were classified according to the concordant direction of regulation between species based on the log-fold change. All DEGs with concordant directionality across the two datasets and with FDR < 0.05 were used for pathway analysis in ShinyGO (v0.76).

**Genomic DNA extraction and HRM-based genotyping.** For genotyping, zebrafish larval tails were excised on ice and placed into TE buffer. Samples were incubated in a thermocycler at 95°C for 10 minutes to initiate tissue lysis. Proteinase K (0.25 mg/mL; A&A Biotechnology, #1019) was then added, and the samples were incubated at 55 °C for a minimum of 4 hours to complete digestion. Proteinase K was subsequently inactivated by heating at 95 °C for 10 minutes. The resulting genomic DNA was used directly for high-resolution melting (HRM) analysis. HRM was performed using qPCR-HS Mix EvaGreen (A&A Biotechnology, #2008HS-1000G) on a LightCycler 96 System (Roche), following the manufacturer's instructions.

**Spontaneous tail flick and body coil behavior.** Spontaneous tail flick and body coil behavior is associated with seizure activity, as described previously (11, 12). 5 hpf, embryos were treated with either DMSO or BAF inhibitor. Following overnight incubation at 28 °C, coiling behavior was recorded under a Leica M60 microscope with a DMC2900 camera every 10 ms for 3 minutes, with a maximum of 12 embryos per plate. Videos were analyzed manually, counting the total and double coils for each larva. A similar analysis was performed on *smarca4* mutant zebrafish. Additionally, a rescue experiment for the spontaneous tail flick and body coil behavior in the *smarca4* line was performed by injecting embryos at the one-cell stage with 200 ng/µL of mouse *Brg1* RNA synthesized from the pIVT/Smarca4 plasmid (gift from Carmen Williams; Addgene plasmid # 122146; http://n2t.net/addgene:122146; RRID:Addgene_122146)). RNA was synthesized using the HiScribe Quick T7 High Yield RNA Synthesis Kit (NEB, #E2050) according to the manufacturer's instructions. Following transcription, samples were treated with DNase I, purified using the Monarch Spin RNA Cleanup Kit (NEB, #T2040), quantified with a NanoDrop, and stored at -80 °C.

**Locomotor behavior.** Locomotor behavior was conducted based on a previously published protocol (12). After 24 or 48 hours of BAF inhibitor incubation, locomotor behavior was assessed using the Zebrabox system (ViewPoint, France). A 24-well plate, with one larva per well, was prepared 24 hours before the behavioral test for habituation. On the day of the experiment, the fish were allowed to acclimate to the behavioral room and zebrabox for at least 10 minutes before starting the test. The 24-well plate was then placed in the Zebrabox, and behavior was recorded for 1 hour in complete darkness. Locomotor activity was categorized as medium (0.5 - 2 cm/sec) or fast swimming (>2 cm/sec). Similar analysis was conducted on the *smarca4* mutant line. Data were analyzed using RStudio (v2021.09.0+351; https://posit.co/).

**RT-qPCR.** Twenty zebrafish larvae or thirty heads were pooled and stored at -20 °C in stayRNA buffer (#038-100, A&A Biotechnology) for RNA preservation until further use. Total RNA was extracted using the Monarch Total RNA Miniprep Kit (NEB, #T2110), following the manufacturer's instructions. cDNA was synthesized using the High-Capacity cDNA Reverse Transcription Kit (ThermoFisher, #4368814) according to the recommended protocol. qPCR was performed with qPCR-HS Mix EvaGreen (A&A Biotechnology, #2008HS-1000G) on a LightCycler 96 system (Roche). The fold change for each gene was calculated using the ΔΔCq method. The primers used for the amplification are the following: **Brg1-related genes:** *klf2a*-F: 5'-AACTGGGACGACTGGAAATGG-3'; *klf2a*-R: 5'-CATCCTTCCACCTGTTCTCCC-3'. *nos1*-F: 5'-GGAGCACGAGACACTTGTGA-3'; *nos1*-R: 5'-CGCCGCACCAAATTTCTCTC-3'. *smarca4*-F: 5'-CGTAAAGAGCTGCCCGAGTA-3'; *smarca4*-R: 5'-GACTCCTGATGCGCTCCTTG-3'. **Proinflammatory-related genes:** *irf1b*-F: 5'-GTGGGTCAACAAGGAGGAGA-3'; *irf1b*-R: 5'-

TGCTTGAACAGACAGGCATC-3'. *il1b*-F: 5'-GCTGGAGATCCAAACGGATA-3'; *il1b*-R: 5'-ATACGCGGTGCTGATAAACC-3'. *il8*-F: 5'-GTCGCTGCATTGAAACAGAA-3'; *il8*-R: 5'-AGGGGTCCAGACAGATCTCC-3'. **Inflammation related to human temporal lobe epilepsy:** *csf1ra*-F: 5'-ACTTTCCAGAACCCATGACG-3'; *csf1ra*-R: 5'-CTCCGACGAAGAATCCAGAG-3'. *c4*-F: 5'-ACAGTGAAGGGAGAGCTGGA-3'; *c4*-R: 5'-GCTCATGGGCTCATCATTTT-3'. *nrros*-F: 5'-AGCCGAAACAGGCTAACTGA-3'; *nrros*-R: 5'-TTGGTGGGAAGTTCTGAAGG-3'. **RNA seq DEGs-related genes:** *gabra6a*-F: 5'-GGCTTCGGCTCTACCTCAGTC-3'; *gabra6a*-R: 5'-GCGCTTCAGACACAGAGTAAC-3'. *slc38a3b*-F: 5'-ATCCTGGCCTTTGCCTTTGT-3'; *slc38a3b*-R: 5'-TTCTGTGTTGGGTTGCGGAG-3'. *caspbl*-F: 5'-GCTGTTTAGGAAGGTTGCCG-3'; *caspbl*-R: 5'-TGGTAAGAGTTTCGCCTGACG-3'. *plp1a*-F: 5'-TCCTTTCTGAAACGCCCTCC-3'; *plp1a*-R: 5'-GCAGTCCATGGAAGTAAACCGTA-3'. **Housekeeping gene for normalization:** *gapdh*-F: 5'-GTGGAGTCTACTGGTGTCTTC-3'; *gapdh*-R: 5'-GTGCAGGAGGCATTGCTTACA-3'.

**Whole-mount immunostaining.** Whole-mount immunostaining was conducted based on a previously established protocol (zfin.org)(12). Zebrafish larvae were fixed in 4% paraformaldehyde (PFA) at 4 °C overnight. Following fixation, samples were washed in PBS and bleached in a freshly prepared solution of 3% $H_2O_2$ and 1% KOH in water. Larvae were permeabilized through sequential washes with PBS containing 0.2% Triton X-100 (#T8787, Sigma-Aldrich), DMSO (#D8418, Sigma-Aldrich), and Tween-20 (#P1379, Sigma-Aldrich). Antigen retrieval was performed using Tris buffer (150 mM, pH 9; #TRS001, Bioshop) at 70 °C for 15 minutes. Blocking was carried out overnight at room temperature in PBS containing 0.2% Triton X-100, 20% DMSO, 0.3 M glycine, and 6% donkey serum. Primary antibody incubation was performed at room temperature over two nights in PBS with 0.2% Tween-20 and 10 µg/mL heparin, followed by extensive washes. Secondary antibody incubation (1:1000, Thermo Fisher) was carried out under the same buffer conditions, protected from light, also over two nights. Larvae were subsequently cleared through a graded glycerol series (30%, 50%, and 70% in PBS) and stored in mounting medium (2% propyl gallate in 90% glycerol with PBS) at 4 °C in the dark. Fluorescence intensity was acquired at the Lightsheet microscope Z1 (Zeiss) with a magnification of 40x. Z-stack acquisition was performed over an approximate range of 100 μm with 50 slices at 2.00 µm intervals. Hoechst$^+$ cell analysis was conducted by applying Otsu thresholding followed by a binary watershed algorithm. Cell numbers were then quantified by particle analysis. The primary antibodies used are the following: vGlut1/2 (1:400, Synaptic System, #135503), GAD65/67 (1:400, abcam, #ab11070), Gephyrin (1:200, BD Transduction Laboratories, #610584), GFAP (1:100, ZIRC, #ZDB-ATB-081002-46), mCherry (1:500, Sicgen, #AB0040-200). Image analyses were performed using Fiji ImageJ (v1.53q, imagej.net/ij/) (13). Fluorescence intensity was measured using the measure stack tool.

**Calcium imaging.** *Tg(HuC:GCaMP5G)* or *Tg(HuC:GCaMP5G)* crossed with *smarca4* mutant line embryos at 8 hpf were treated daily with 0.003% 1-phenyl 2-thiourea (PTU). Imaging was conducted based on a previously published protocol (12). Larvae at 119 hpf were incubated for 30 minutes in 300 μM Pancuronium bromide (#P1918, Sigma-Aldrich) to inhibit movements without affecting brain activity. Then, larvae were exposed to a subthreshold dose of PTZ (1.25 mM) for 3 minutes. Calcium influx was imaged at the Lightsheet microscope (Zeiss) with a magnification of 40x. 3D-time-lapse images of neuronal activity were taken for the optic tectum, with an approximate range of 100 μm recorded for 3 minutes. For neuronal activity image analysis, first-drift correction was performed in Fiji ImageJ (v1.53q, imagej.net/ij/) (13) and then the fluorescence intensity was calculated using the measure stack tool. Changes in intracellular calcium levels were quantified by the relative fluorescence change (deltaF). deltaF was calculated using the following formula: $\Delta F = (F_{t+1} - F_t) / F_{min}$, where $F_t$ and $F_{t+1}$ are the mean fluorescence intensities at frames *t* and *t+1*, respectively, and $F_{min}$ is the minimum fluorescence value across the entire dataset.

**Acridine orange staining.** Zebrafish larvae previously treated with either DMSO or the BAF inhibitor for 24 or 48 hours were incubated in an acridine orange solution (0.3 mg/mL in E3 medium) for 30 minutes in the dark. After incubation, larvae were sedated with tricaine and imaged using light sheet microscope (Zeiss) with a 40x magnification. To block the skin pigmentation, embryos were treated daily with 0.003% PTU till the day of image acquisition. Fluorescent cells were counted manually with the multipoint tool in Fiji ImageJ (v1.53q; imagej.net/ij/) (13).

**Non-invasive local field potential (LFP) recordings.** 96 hpf zebrafish larvae were treated overnight with either DMSO or BAF inhibitor. Following treatment, 115-125 hpf larvae were exposed to a subthreshold concentration of PTZ (6 mM) for 10 minutes. Non-invasive local field potential (LFP) recordings were carried out as previously described (14, 15). Larvae were immobilized in 2% low-melting-point agarose, whereafter electrical signals were measured for 10 minutes from the optic tectum (midbrain) via a glass electrode, filled with artificial cerebrospinal fluid (124 mM NaCl, 2 mM KCl, 2 mM $MgSO_4$, 2 mM $CaCl_2$, 1.25 mM $KH_2PO_4$, 26 mM $NaHCO_3$, and 10 mM glucose). Clampfit 10.2 software (Molecular Devices Corporation) was used to visualize epileptiform events. LFP recordings were analyzed using Welch's method for power spectral density (PSD), applying 100 ms segments with Hamming windowing and 80% overlap between segments. For each recording, the average power spectral density was calculated across consecutive 10 Hz frequency bands.

**Mass spectrometry.** 96 hpf, *smarca4* mutant zebrafish were euthanized, and their heads were dissected on ice. The heads were immediately snap-frozen in liquid nitrogen for downstream analysis, while the tails were used for HRM-based genotyping. Following genotyping, heads from five individuals per genotype were pooled to create five biological replicates. Tissue lysis and protein extraction were done according to the Sample Preparation by Easy Extraction and Digestion (SPEED) protocol (16). Tissues were solubilized in concentrated Trifluoroacetic Acid (TFA; Cat: T6508; Sigma Aldrich) (sample /TFA 1:4 (v/v)) and incubated for 10 min at room temperature and sonicated in a water bath (~30 s). Next, samples were neutralized with 2 M Tris-Base buffer using 10x volume of TFA and further incubated at 95 °C for 5 min after adding Tris(2-carboxyethyl)phosphine (final concentration 10 mM) and 2-chloroacetamide (final concentration 40 mM). Protein concentrations were determined by turbidity measurements at 360 nm, adjusted to the same concentration using a sample dilution buffer (2M TrisBase/TFA 10:1 (v/v)), and then diluted 1:3 with water. Protein digestion was carried out overnight at 37 °C using sequencing-grade modified trypsin (Promega) at a protein/enzyme ratio of 20:1. The digestion was terminated by the addition of TFA to 2% final concentration. The resulting peptides were labelled with TMT using an on-column TMT labeling protocol (17). Stage tips were packed with three punches of C18 mesh (Affinisep) with a 16-gauge blunt-end needle. Resin was conditioned with 150 µl methanol (MeOH), followed by 100 µl 50% acetonitrile (ACN)/0.1% formic acid (FA), and equilibrated with 150 µl 0.1% FA twice. The digest was loaded by spinning at 1200 × g until the entire digest passed through. The bound peptides were washed twice with 150 µl 0.1% FA. 200 µl of TMT reagent in 50 mM phosphate buffer pH 8 (TMT reagent was resuspended in 2 µl 100% ACN and diluted with 200 µl of 50 mM phosphate buffer pH 8) was loaded over the C18 resin at 300 × g until the entire solution had passed through. The buffer and residual TMT were washed away three times with 150 µl 0.1% FA. Peptides were eluted with 60 µl 60% ACN/0.1% FA. An equal volume of each sample was pooled in one microcentrifuge tube and dried in SpeedVac. Peptides in the compiled sample were separated into 8 fractions using the Pierce High pH Reversed-Phase Peptide Fractionation Kit (Thermo Fisher Scientific). Prior to LC-MS/MS measurements, the peptide fractions were resuspended in 0.1% TFA, 2% acetonitrile in water.
Chromatographic separation was performed on an Easy-Spray Acclaim PepMap column 50 cm long × 75 µm inner diameter (Thermo Fisher Scientific) at 55 °C by applying a 120 min acetonitrile gradients in 0.1% aqueous formic acid at a flow rate of 300 nl/min. An UltiMate 3000 nano-LC system was coupled to a Q Exactive HF-X mass spectrometer via an easy-spray source (all Thermo Fisher Scientific). The Q Exactive HF-X was operated in TMT mode with survey scans acquired at a resolution of 60,000 at m/z 200. Up to 15 of the most abundant isotope patterns with charges 2-5 from the survey scan were selected with an isolation window of 0.7 m/z and fragmented by higher-energy collision dissociation (HCD) with normalized collision energies of 32, while the dynamic exclusion was set to 35 s. The maximum ion injection times for the survey scan and the MS/MS scans (acquired with a resolution of 45,000 at m/z 200) were 50 and 120 ms, respectively. The ion target value for MS was set to 3e6 and for MS/MS to 1e5, and the minimum AGC target was set to 1e3.
The data were processed with MaxQuant v. 1.6.17.0 (18), and the peptides were identified from the MS/MS spectra searched against Uniprot zebrafish reference proteome (UP000000437) using

the built-in Andromeda search engine. Reporter ion MS2-based quantification was applied with reporter mass tolerance = 0.003 Da and min. reporter PIF = 0.75. Cysteine carbamidomethylation was set as a fixed modification, and methionine oxidation, asparagine and glutamine deamidation as well as protein N-terminal acetylation were set as variable modifications. For in silico digests of the reference proteome, cleavages of arginine or lysine followed by any amino acid were allowed (trypsin/P), and up to two missed cleavages were allowed. The FDR was set to 0.01 for peptides, proteins, and sites. Match between runs was enabled. Other parameters were used as pre-set in the software.

Reporter intensity corrected values for protein groups were loaded into Perseus v. 1.6.10 (19). Standard filtering steps were applied to clean up the dataset: reverse (matched to decoy database), only identified by site, and potential contaminant (from a list of commonly occurring contaminants included in MaxQuant) protein groups were removed. Reporter intensity corrected values were log2 transformed, and protein groups with values across all samples were kept. The values were normalized by median subtraction within TMT channels. Student's t-test (2-sided, permutation-based FDR = 0.001, S0 = 0.1, n = 5) was performed to determine proteins differentially regulated between the two sample groups. The data were exported from Perseus and formatted to its final form in Microsoft Excel 2016.

The differentially regulated proteins were used as input for pathway analysis. Functional enrichment analysis was performed using ShinyGO (v0.76) (7) to identify significantly affected pathways across various ontology categories, including GO:BP, GO:CC, and GO:MF. Additionally, enrichment was assessed using the Zebrafish disease, KEGG, anatomy, and phenotype databases. For the KEGG analysis, gene IDs were converted from zebrafish to human to improve annotation quality and reveal additional potential pathways.

**Statistical analysis.** All statistical analyses were performed using GraphPad Prism 10 software. Exact sample sizes for each experiment are indicated in the figure legends. For data following a normal distribution, statistical comparisons were performed using Student's *t*-test, one-way ANOVA, or two-way ANOVA, followed by appropriate post hoc tests, including Dunnett's, Tukey's, or Dunn's test for multiple comparisons. In cases of unequal variance, Welch's *t*-test and Brown-Forsythe ANOVA followed by Dunnett's T3 multiple comparisons test were used. For data not conforming to a normal distribution, non-parametric tests such as the Mann-Whitney *U* test or the Kruskal-Wallis test followed by appropriate post hoc tests were used. Data are presented as mean ± standard error of the mean (SEM). A *p*-value of less than 0.05 was considered statistically significant. Unless otherwise stated, all statistical tests were two-sided.

**Figures**

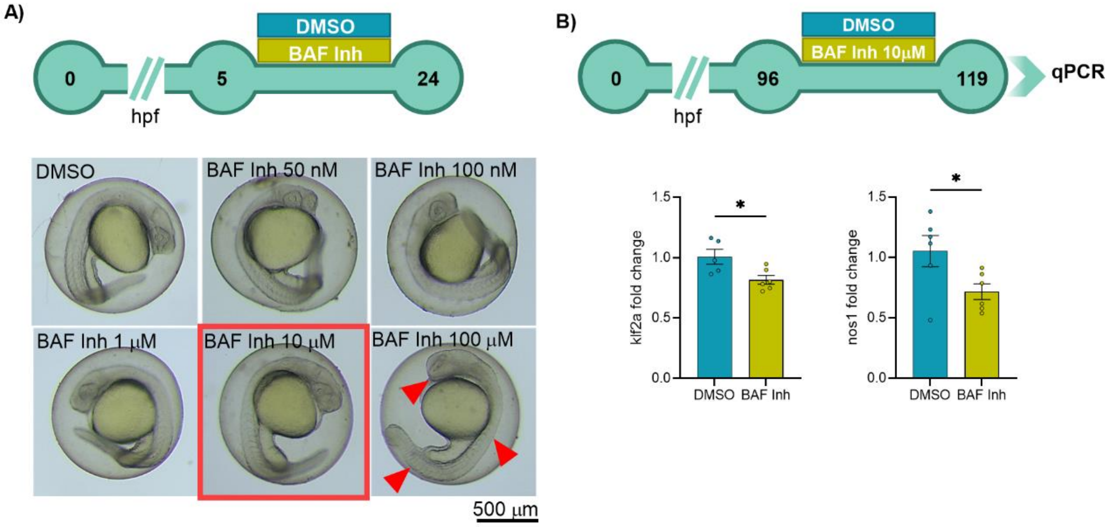


**Figure S1. Functional validation of BAF complex inhibitor in zebrafish model. (A)** Experimental timeline. Zebrafish larvae were treated at 5 hours post-fertilization (hpf) with various concentrations of either DMSO or a BAF complex inhibitor (BAF Inh.; 50 nM, 100 nM, 1 µM, 10 µM, and 100 µM). Following overnight treatment, larvae were examined for phenotypic abnormalities. The highest concentration that did not alter larval morphology was selected for further experiments (10 µM). **(B)** Experimental timeline. Zebrafish larvae at 96 hpf were treated with 10 µM BAF Inh or 0.02% DMSO as a control. To validate the inhibitor's functionality, the expression of known Brg1-regulated genes was assessed: *klf2a* ($t(9) = 2.785$, $p = 0.0212$), *nos1* ($t(10) = 2.332$, $p = 0.0419$) ($n_{DMSO} = 5\text{-}6$, $n_{BAF\ Inh} = 6$ replicates; each replicate consisted of 20 larvae). Data are presented as mean ± standard error of the mean (SEM), $^*p < 0.05$.

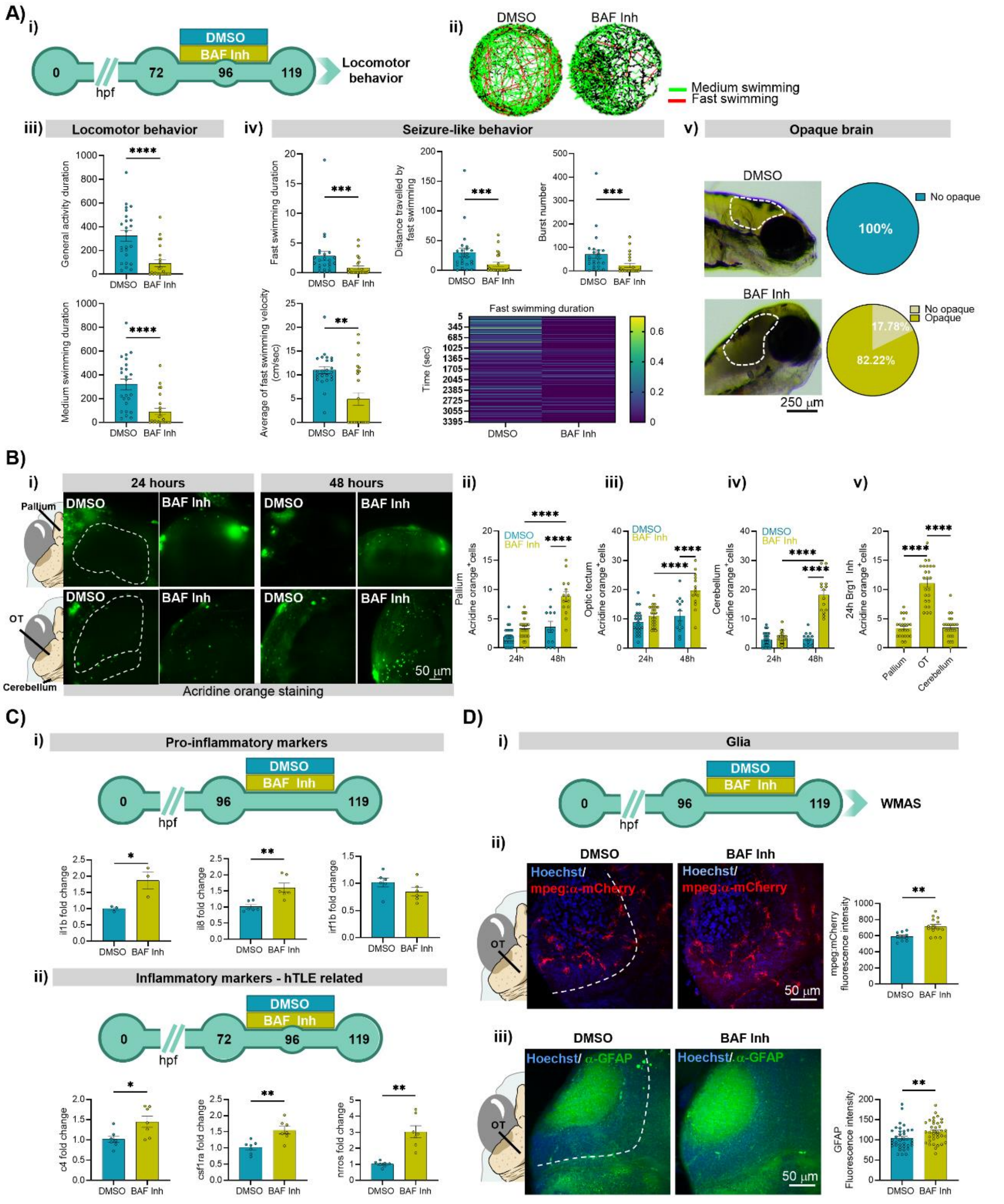

**Figure S2. BAF complex inhibition induces inflammation, cellular death and activation of glia in the zebrafish larvae brain. A) i)** Experimental timeline. Zebrafish larvae were treated at 72 hours post-fertilization (hpf) with either DMSO or BAF complex inhibitor (BAF Inh, 10 µM). At 119 hpf, larvae were tested for locomotor behavior. **ii)** Representative locomotor tracks of larvae treated with DMSO or BAF Inh. **iii)** General locomotor activity: Duration of general activity (Mann-Whitney test, U = 88, $p < 0.0001$) and duration of medium swimming (Mann-Whitney test, U = 88, $p < 0.0001$) after BAF Inh. Data are presented as mean ± standard error of the mean (SEM). **iv)** Seizure-like behavior parameters: Fast swimming duration (Mann-Whitney test, U = 112.5, $p = 0.0002$), distance traveled with fast swimming (Mann-Whitney test, U = 119, $p = 0.0003$), number of burst episodes

(Mann-Whitney test, U = 115, $p = 0.0002$), average velocity during fast swimming (Mann-Whitney test, U = 141, $p = 0.0018$), and a heatmap illustrating fast swimming duration over time ($n_{DMSO}$ = 24, $n_{BAFInh}$ = 24). Data are presented as mean ± SEM. **v)** Representative image of larvae showing brain opacity following 48-hour treatment with the BAF Inh. Pie charts showing the percentage of larvae with opaque brains after 48 hours of treatment with either DMSO or the BAF Inh ($n_{DMSO}$ = 41, $n_{BAFInh}$ = 45). **(B) i)** Zebrafish larvae were treated at 96 hpf with either DMSO or BAF Inh or at 72 hpf for 48 hours. At 119 hpf, acridine orange staining was employed to assess neuronal death in pallium, optic tectum (OT), and cerebellum. **ii)** Quantification of fluorescent cells number in the pallium after 24 or 48 hours of treatment with the BAF Inh (two-way ANOVA: Time effect, F (1, 70) = 37.21, $p < 0.0001$; treatment effect, F (1, 70) = 32.17, $p < 0.0001$; Time x Treatment interaction, F (1, 70) = 10.52, $p = 0.0018$; Tukey post hoc test), ($n_{DMSO\_24h}$ = 24, $n_{DMSO\_48h}$ = 12, $n_{BAFInh\_24h}$ = 24, $n_{BAFInh\_48h}$ = 14). Data are presented as mean ± SEM. **iii)** Quantification of fluorescent cells number in the OT after 24 or 48 hours of treatment with the BAF Inh (two-way ANOVA: Time effect, F (1, 71) = 23.32, $p < 0.0001$; treatment effect, F (1, 71) = 23.37, $p < 0.0001$; Time x Treatment interaction, F (1, 71) = 8.213, $p = 0.0055$; Tukey post hoc test), ($n_{DMSO\_24h}$ = 24, $n_{DMSO\_48h}$ = 13, $n_{BAFInh\_24h}$ = 24, $n_{BAFInh\_48h}$ = 14). Data are presented as mean ± SEM. **iv)** Quantification of fluorescent cells number in the cerebellum after 24 or 48 hours of treatment with the BAF Inh (two-way ANOVA: Time effect, F (1, 71) = 86.78, $p < 0.0001$; treatment effect, F (1, 71) = 93.97, $p < 0.0001$; Time x Treatment interaction, F (1, 71) = 81.37, $p < 0.0001$; Tukey post hoc test), ($n_{DMSO\_24h}$ = 24, $n_{DMSO\_48h}$ = 13, $n_{BAFInh\_24h}$ = 24, $n_{BAFInh\_48h}$ = 14). Data are presented as mean ± SEM. **v)** Comparison between the three brain regions after 24 hours BAF Inh treatment (Brown-Forsythe ANOVA: F (2, 49.59) = 75.33, $p < 0.0001$; Dunnett's T3 multiple comparisons test), ($n_{BAFInh\_each\_brain\_regions}$ = 24). Data are presented as mean ± SEM. **(C) i)** RT-qPCR of proinflammatory markers in zebrafish larvae after 24 hours BAF Inh treatment (*il1b*: t(4) = 3.23, $p = 0.0321$; *il8*: t(10) = 3.82, $p = 0.0034$; *irf1b*: t(10) = 1.51, $p = 0.1629$), ($n_{DMSO}$ = 3-6, $n_{BAFInh}$ = 3-6; each replicate consisted of 30 larvae). Data are presented as mean ± SEM. **ii)** RT-qPCR of inflammatory markers in zebrafish larvae after 48 hours BAF Inh treatment (*c4*: t(12) = 2.73, $p = 0.0184$; *csf1ra*: t(12) = 3.85; $p = 0.0023$; *nrros*: *t*(6.376) = 5.300, $p = 0.0015$, Welch's correction), ($n_{DMSO}$ = 7, $n_{BAFInh}$ = 7; each replicate consisted of 30 larvae heads). Data are presented as mean ± SEM. **(D) i)** Experimental timeline. Zebrafish larvae were treated at 96 hpf with either DMSO or BAF Inh. At 119 hpf, larvae were sacrificed and used for whole-mount immunofluorescence (IF) staining for glia markers (microglia: mCherry driven by mpeg promoter; astrocytes: GFAP). **ii)** Representative images Z-projection (maximum intensity) showing mpeg staining in the OT of larvae treated with DMSO or BAF Inh. Quantification of mCherry IF in the OT of DMSO and BAF Inh treated zebrafish (t(22) = 3.45, $p = 0.0023$; $n_{DMSO}$ = 9, $n_{BAFInh}$ = 15). Data are presented as mean ± SEM. **iii)** Representative Z-projection (maximum intensity) images showing GFAP IF staining in the OT of larvae treated with DMSO or BAF Inh. Quantification of GFAP IF intensity in the OT of DMSO and BAF Inh treated zebrafish (Mann-Whitney test, U = 421, $p = 0.0065$; $n_{DMSO}$ = 36, $n_{BAFInh}$ = 37). Data are presented as mean ± SEM. *$p < 0.05$, **$p < 0.01$, ***$p < 0.001$, ****$p < 0.0001$.

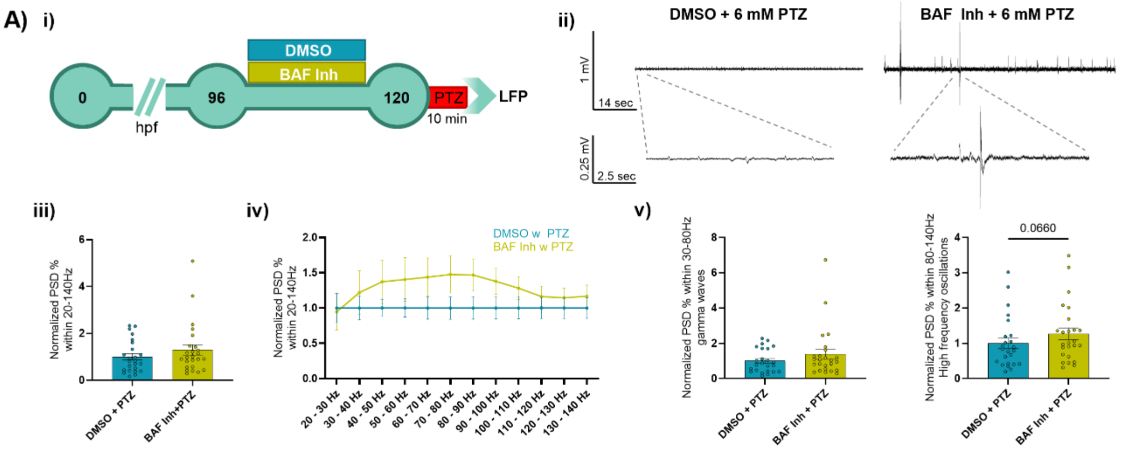


**Figure S3. Inhibition of the BAF complex tends to increase brain activity following a subthreshold dose of PTZ. (A) i)** Experimental timeline. Zebrafish larvae were treated at 96 hours post-fertilization (hpf) with either DMSO or BAF complex inhibitor (BAF Inh, 10 µM). At 120 hpf, larvae underwent non-invasive local field potential (LFP) recordings after exposure to a subthreshold dose of pentylenetetrazol (PTZ, 6 mM). **ii)** Representative LFP recordings from larvae treated with either DMSO or the BAF Inh following exposure to a subthreshold dose of PTZ. **iii)** Normalized power spectral density (PSD) after BAF Inh (Mann-Whitney test, U = 251, $p$ = 0.3352). **iv)** PSD distribution across frequency bands from 20-140 Hz (two-way ANOVA: Hz effect, F (2.244, 105.5) = 1.737, $p$ = 0.1770; treatment effect, F (1, 47) = 1.219, $p$ = 0.2752; Hz x treatment effect, F (11, 517) = 1.737, $p$ = 0.0625). **v)** Normalized PSD within the frequency of 30–80 Hz (Mann-Whitney test, U = 255, $p$ = 0.1884, One-tailed), and 80–140 Hz (Mann-Whitney test, U = 224, $p$ = 0.0660, One-tailed) ($n_{DMSO\_PTZ}$ = 24, $n_{BAFInh\_PTZ}$ = 25). Data are presented as mean ± standard error of the mean (SEM).

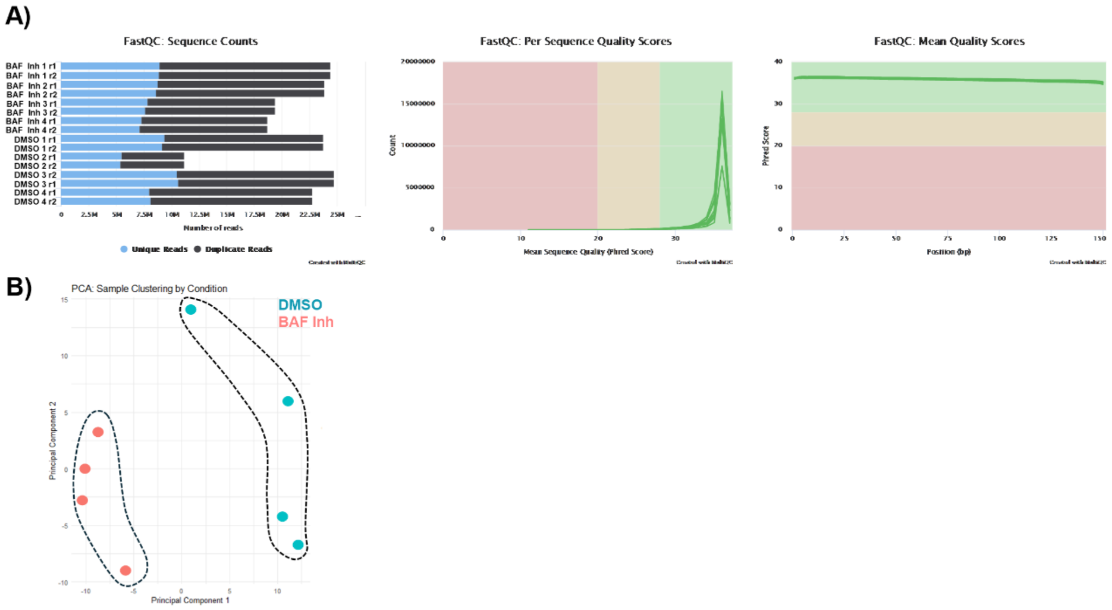


**Figure S4. Quality control assessment and sample clustering of RNA-seq data. (A)** Sequencing quality control. Left: Sequence counts for each sample, showing the number of unique and duplicate reads (FastQC). Middle: Per-sequence quality score distribution, indicating high overall read quality. Right: Mean quality scores per base position across all reads, demonstrating consistently high-quality sequences. **(B)** Principal component analysis (PCA) of normalized gene expression profiles. Samples cluster distinctly by experimental condition, with BAF inhibitor (red) and DMSO (blue) groups forming separate clusters, indicating clear separation.

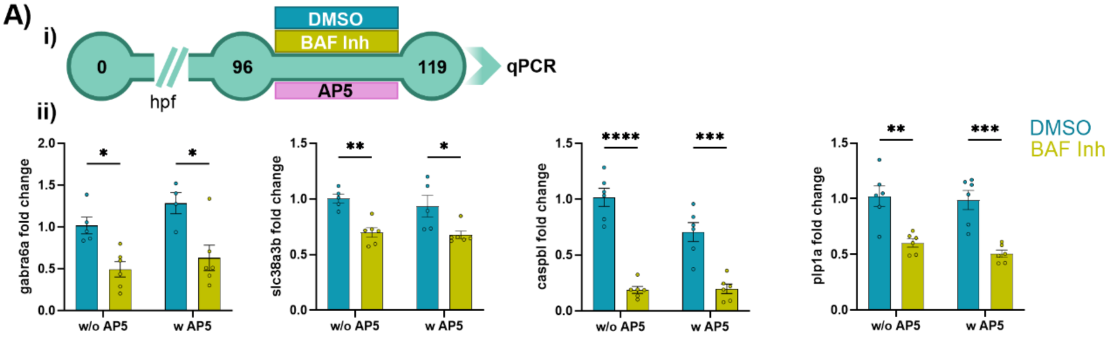


**Figure S5. The molecular impairments are specifically caused by BAF inhibition, rather than by seizure development. (A) i)** Experimental timeline. Zebrafish larvae were treated at 96 hours post-fertilization (hpf) with either DMSO or a BAF complex inhibitor (BAF Inh, 10 μM), along with (2R)-amino-5-phosphonovaleric acid (AP5, 100 μM) to block seizure development. At 119 hpf, the larvae were sacrificed and subjected to RT-qPCR to measure the expression of genes that were found to be deregulated in the RNA sequencing experiment. **ii)** mRNA level of *gabra6a* (two-way ANOVA: treatment effect, $F(1, 17) = 22.59$, $p = 0.0002$; Tukey post hoc test), *slc38a3b* (two-way ANOVA: treatment effect, $F(1, 18) = 24.88$, $p < 0.0001$; Tukey post hoc test), *caspbl* (two-way ANOVA: treatment effect, $F(1, 20) = 106.9$, $p < 0.0001$; AP5 effect, $F(1, 20) = 5.448$, $p = 0.0301$; Treatment vs. AP5 interaction, $F(1, 20) = 6.152$, $p = 0.0221$; Tukey post hoc test) and *plp1a* (two-way ANOVA: treatment effect, $F(1, 20) = 44.12$, $p < 0.0001$; Tukey post hoc test), ($n_{DMSO} = 5\text{-}6$, $n_{DMSO\_AP5} = 4\text{-}6$ $n_{BAFInh} = 6$, $n_{BAFInh\_AP5} = 6$ replicates, each replicate consists in 30 larvae heads). Data are presented as mean ± standard error of the mean (SEM). *$p < 0.05$, **$p < 0.01$, ***$p < 0.001$, ****$p < 0.0001$.

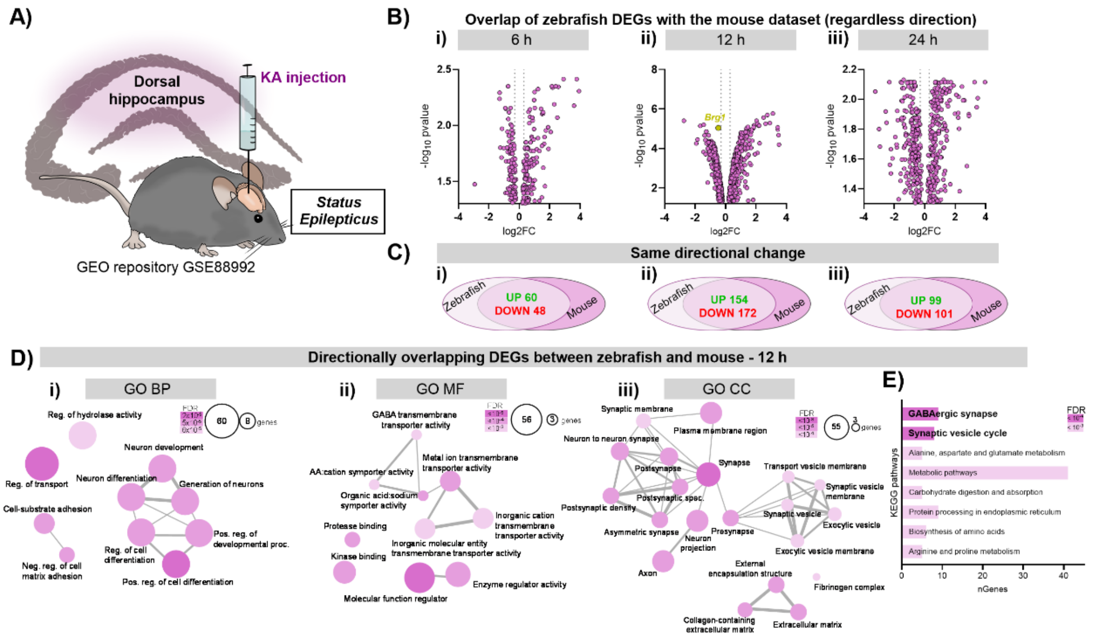


**Figure S6. Comparative transcriptomic analysis of zebrafish BAF inhibitor-treated larvae and a mouse kainic acid-induced epilepsy model. (A)** Experimental design adapted from Kalozoumi et al. (2018). C57BL/6J mice were injected into the dorsal hippocampus with kainic acid (KA) or saline to induce the *status epilepticus* (SE) and were sacrificed at 6, 12, and 24 hours post-injection (GSE88992). **(B) i-iii)** Volcano plots illustrating the shared differentially expressed genes (DEGs) between KA-injected mice and BAF inhibitor-treated zebrafish. **(C) i-iii)** Venn's diagrams showing the number of genes deregulated in the same direction in the mouse and zebrafish datasets. **(D) i-iii)** Gene Ontology (GO) network analysis of shared DEGs with concordant expression changes at the 12-hour time point, including Biological Process (GO BP), Molecular Function (GO MF), and Cellular Component (GO CC) categories. **iv)** Kyoto Encyclopedia of Genes and Genomes (KEGG) pathway enrichment analysis of shared DEGs with concordant expression changes at the 12-hour time point.

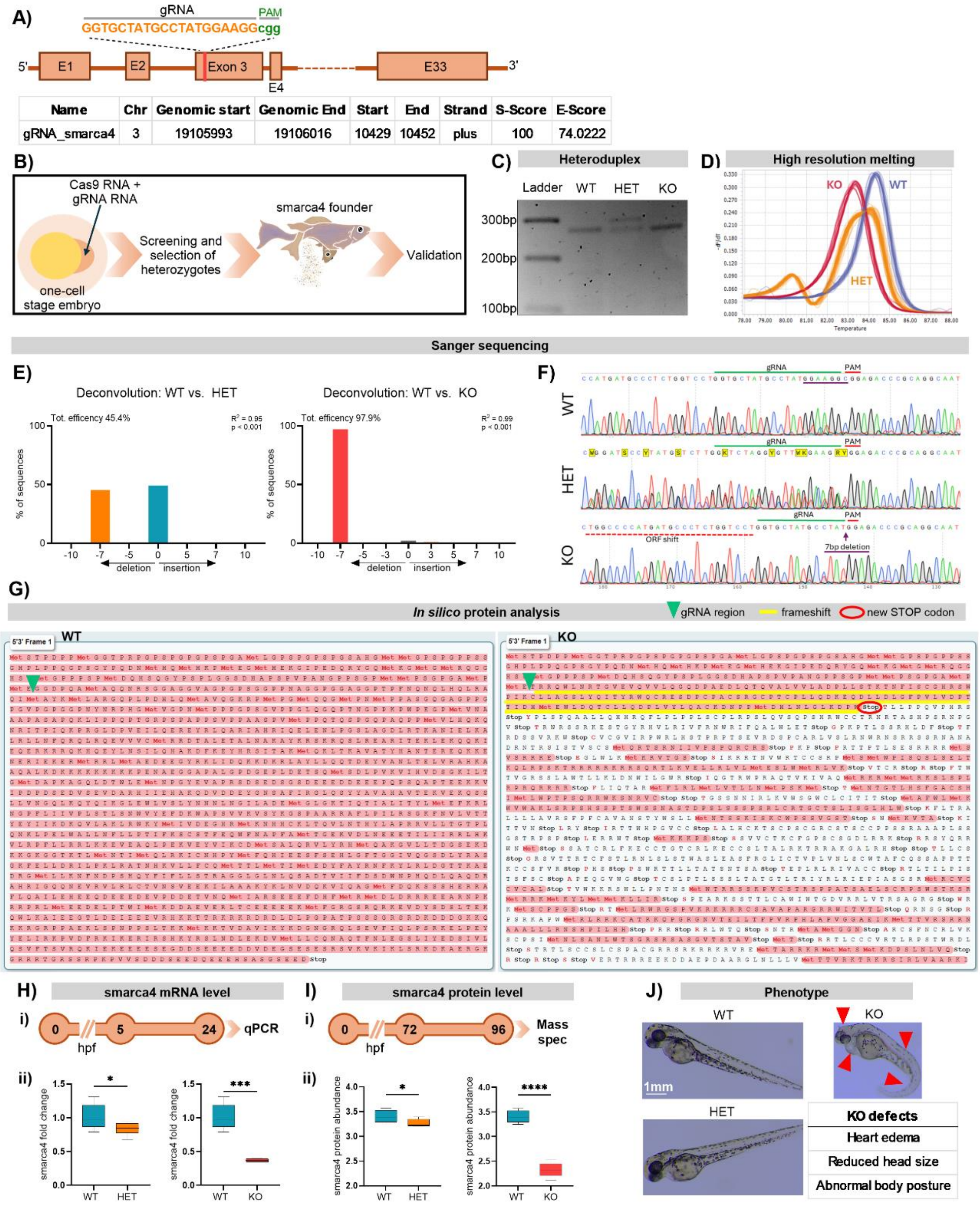


**Figure S7. Generation and validation of CRISPR/Cas9-mediated *smarca4* mutant line. (A)** A gRNA was selected for CRISPR/Cas9 editing of exon 3 of the *smarca4* gene. The table shows the gRNA's genomic location, specificity, and efficiency scores. **(B)** Schematic representation of the generation of the *smarca4* mutant line using the CRISPR/Cas9 system. One-cell stage embryos were injected with gRNA and Cas9 mRNA. Embryos were screened for successful mutations and raised to adulthood. Founder fish were then crossed with wild-type (WT; AB x TL strain) fish to establish and validate the mutant line. **(C)** Gel electrophoresis showing heteroduplex formation in heterozygote (HET) sample. **(D)** High-resolution melting (HRM) analysis was used for genotyping WT, HET, and knockout (KO) larvae. WT samples displayed a single, sharp melting curve, characteristic of homoduplex DNA. HET samples exhibited broader, shifted melting curves due to the presence of both homoduplex and heteroduplex DNA. KO samples, carrying homozygous mutations, showed distinct melting profiles different from both WT and HET. **(E)** Sanger sequencing results for WT, HET, and KO. Deconvolution of sequencing traces showed that HET samples contained ~50% WT sequence and ~50% sequence with a 7 bp deletion. KO samples showed ~98% of sequences with the 7 bp deletion. **(F)** Chromatogram comparison of WT, HET, and KO. The HET chromatogram showed overlapping peaks due to a mixture of WT and mutant sequences, while the KO chromatogram clearly displayed a 7 bp deletion in the gRNA target region. **(G)** *In silico* analysis of predicted proteins resulting from the 7 bp deletion in the *smarca4* gene. The WT protein sequence is shown on the left, and the KO sequence on the right. The gRNA target region is marked with a green arrow. In the KO sequence, a frameshift (highlighted in yellow) leads to a premature stop codon (circled). **(H) i)** *smarca4* mRNA levels were measured by qPCR in 24 hours post-fertilization (hpf) *smarca4* mutant embryos. **ii)** Fold change in *smarca4* mRNA expression (WT vs. HET: $t(9) = 1.980$, $p = 0.0396$, One-tailed; WT vs. KO: Welch's *t*-test, $t(4.12) = 7.458$, $p = 0.0008$, One-tailed). Data are presented as box-and-whisker plots showing minimum to maximum values. **(I) i)** *smarca4* protein level was measured by mass spectrometry in 96 hours post-fertilization (hpf) *smarca4* mutant embryos. **ii)** Fold change in *smarca4* protein levels (WT vs. HET: $t(8) = 2.041$, $p = 0.0378$, One-tailed; WT vs. KO: $t(8) = 12.50$, $p < 0.0001$, One-tailed). Data are presented as box-and-whisker plots showing minimum to maximum values. **(J)** Phenotypic characterization of the *smarca4* mutant line. No evident differences were observed between WT and HET. KO embryos exhibited reduced eye development, abnormal body posture, and cardiac edema (indicated by red arrows). $^{*}p < 0.05$, $^{***}p < 0.001$, $^{****}p < 0.0001$.

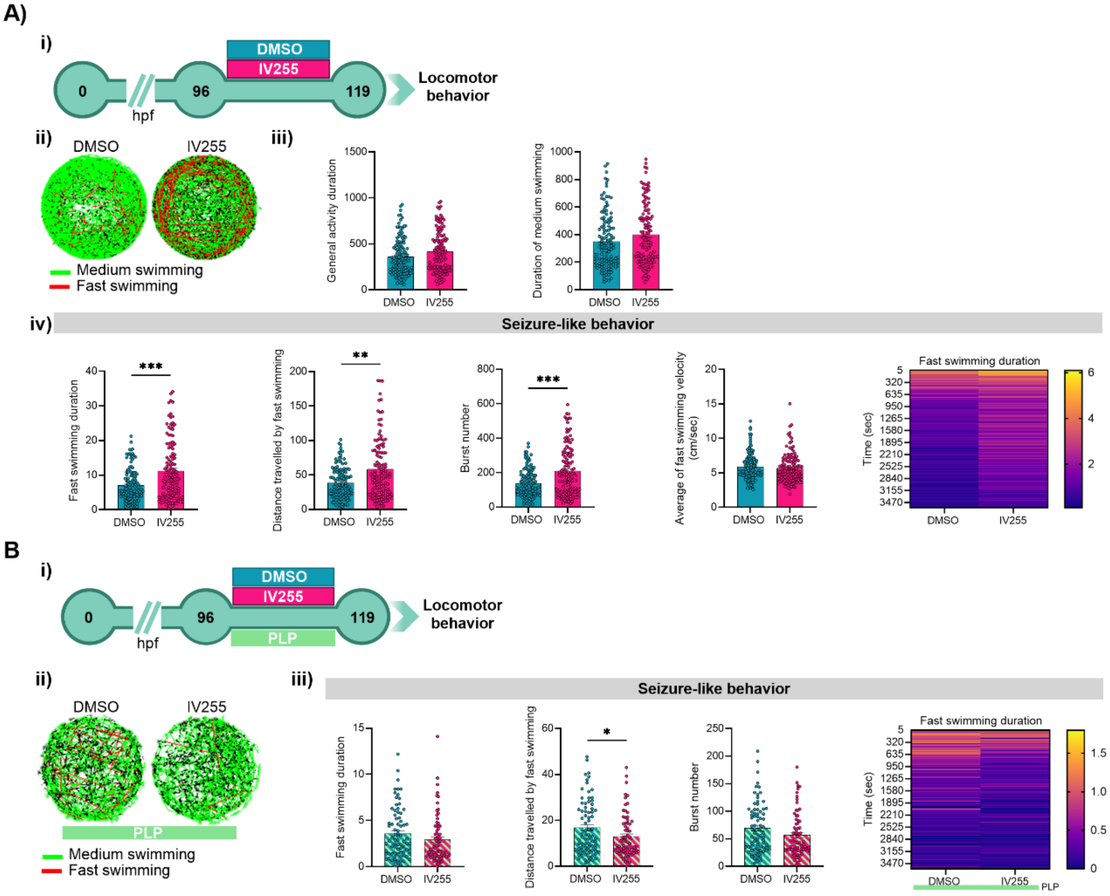


**Figure S8. The specific Brg1 inhibitor IV255 replicated the effects observed with the BAF complex inhibitor and with *smarca4* zebrafish line. (A) i)** Experimental timeline. Zebrafish larvae were treated with IV255 (10 µM) at 96 hours post-fertilization (hpf) and incubated overnight. Locomotor behavior was subsequently assessed. **ii)** Representative locomotor tracks of larvae after IV255 and DMSO treatment. Medium swimming track (0.5–2 cm/sec) is shown in green, while fast swimming track (>2 cm/sec) is shown in red. **iii)** General locomotor activity: Duration of general activity (Mann-Whitney test, U = 6707, $p$ = 0.0532), and duration of medium swimming (Mann-Whitney test, U = 6755, $p$ = 0.0646). Data are presented as mean ± standard error of the mean (SEM). **iv)** Seizure-like behavior parameters: Fast swimming duration (Mann-Whitney test, U = 5753, $p$ = 0.0003), distance traveled with fast swimming (Mann-Whitney test, U = 5972, $p$ = 0.0013), number of burst episodes (Mann-Whitney test, U = 5776, $p$ = 0.0004), average velocity during fast swimming (Mann-Whitney test, U = 7035, $p$ = 0.1744), and a heatmap illustrating fast swimming duration over time ($n_{DMSO}$ = 126, $n_{IV255}$ = 124). Data are presented as mean ± SEM. **(B) i)** Experimental timeline. Zebrafish larvae were treated with IV255 with pyridoxal phosphate (PLP, 100 µM) at 96 hpf and incubated overnight. Locomotor behavior was subsequently assessed. **ii)** Representative locomotor tracks of larvae treated with PLP. Medium swimming track (0.5–2 cm/sec) is shown in green, while fast swimming track (>2 cm/sec) is shown in red. **iii)** Seizure-like behavior parameters: Fast swimming duration (Mann-Whitney test, U = 2373, $p$ = 0.0790), distance traveled with fast swimming (Mann-Whitney test, U = 2317, $p$ = 0.0495), number of burst episodes (Mann-Whitney test, U = 2379, $p$ = 0.0830) and a heatmap illustrating fast swimming duration over time ($n_{DMSO\_PLP}$ = 79, $n_{IV255\_PLP}$ = 72). Data are presented as mean ± SEM. *$p$ < 0.05, **$p$ < 0.01, ***$p$ < 0.001.
